# Time-resolved transcriptomic profiling of senescence-associated secretory phenotype (SASP) in multiple senescent cell subtypes

**DOI:** 10.1101/2022.06.27.497690

**Authors:** Nurhanani Razali, Yohsuke Moriyama, Yatzu Chiu, Kojiro Suda, Keiko Kono

## Abstract

Cellular senescence, irreversible cell cycle arrest, is induced by various triggers including telomere shortening, oncogene activation, and DNA damage. Senescent cells exhibit the senescence-associated secretory phenotype (SASP), a pathological feature that contributes to organismal aging. We previously showed that transient plasma membrane damage (PMD) induces a novel subtype of cellular senescence (PMDS) accompanied by SASP, but the overall expression profiles of SASP during PMDS induction was unknown. Here, using mRNA-seq, qPCR, and bioinformatics, we revealed the time-resolved SASP transcriptomic profile in PMDS in comparison with calcium influx-induced senescence, DNA damage response-induced senescence, and replicative senescence. Although the expression of SASP factors was postulated to increase steadily during senescence, we counterintuitively found that the variety of SASP peaks in early PMDS. The pathway comparison analyses and Ingenuity Pathway Analysis suggest that, in early PMDS, wound-healing SASP factors, namely *Il-6*, *Mmp1*, and *Mmp3,* inhibit the GPVI collagen signaling pathway, which in turn further upregulates the same SASP factors, forming a feedback loop. At late senescence, common SASP factors including *Il-6* and *Ccl*2 are upregulated in all senescent cell subtypes. Thus, SASP is diverse at early senescence and becomes relatively uniform at late senescence. Diverse SASP may contribute to senescent cell subtype-specific paracrine/autocrine functions in vivo.

## INTRODUCTION

Cellular senescence is an essentially irreversible cell cycle arrest that contributes to physiological and pathological processes in vivo, including organismal aging, development, wound healing, and cancer progression (1, 2). Removal of senescent cells ameliorates various age-associated disorders, including cancer, atherosclerosis, and neurodegenerative disorders (2, 3). The three best-described triggers of cellular senescence are telomere shortening, DNA damage, and oncogene activation. The central mechanisms of cellular senescence are the activation of the DNA damage response (DDR) and the upregulation of p53/CDKN1A (p21) and CDKN2A (p16)/pRB pathways (4), frequently associated with senescence features including increased senescence-associated-β-galactosidase (SA-β-gal) activity, oxidative damage, upregulation of anti-apoptotic proteins, and senescence-associated heterochromatin foci (SAHF) (5–7). However, these senescence hallmarks are not uniformly identified in all senescent cell subtypes, raising the possibility that the functions of each senescent subtypes might also be diverse.

In vivo functions of senescent cells are frequently mediated by the senescence-associated secretory phenotype (SASP), namely pro-inflammatory cytokines, chemokines, matrix metalloproteinases (MMP), growth factors, and angiogenic factors secreted by senescent cells (8). SASP can have both beneficial and deleterious effects on the recipient cells. In a beneficial context, the SASP contributes to the recruitment of immune cells that would clear senescent cells and promotes tissue remodeling, wound healing, and tumour suppression (9). In contrast, prolonged SASP exposure can induce detrimental consequences including tissue dysfunction, tumour promotion, and paracrine senescence (10). Thus, SASP is one of the critical factors determining the pathophysiological significance of senescent cells.

We previously reported that the transient plasma membrane damage induced cellular senescence (PMDS) phenotypes over the subsequent 8 – 16 days. PMDS is associated with the upregulation of SASP factors, namely *Il-6* and *Ccl*2, analogous to DNA damage-dependent and replicative senescence (RS) (11). Moreover, we proposed that Ca^2+^ influx after PMD may mediate the induction of p53 and induce DDR-independent premature senescence (calcium influx-induced senescence: CIS) (11). However, the potential in vivo functions of PMDS cells are unknown. Here, we conducted a comprehensive characterization of SASP components and found that transient upregulation of wound healing SASP factors, *Il-6*, *Mmp1,* and *Mmp3,* at the early phase of PMDS potentially modulates the inflammation-mediated injury/repair response. In addition, all senescence inducers tested here, plasma membrane damage, DNA damage response-induced senescence (DDRS), and CIS, induced the persistent upregulation of common SASP factors, *Il-6* and *Ccl2,* at the later phase. These findings suggest that 1) SASP mediates both universal and senescence subtype-specific cellular functions and 2) the SASP-mediates diverse functions in the early phase and relatively uniform functions in the later phase. This work will serve as a basis for further studies to understand how senescent cells can drive diverse pathophysiological functions in vivo.

## MATERIAL AND METHODS

### Cell culture

The human embryonic fibroblast WI-38 was acquired from RIKEN Bioresource Research Center (BRC) and cultured in a humidified incubator at 37 °C and 5% CO_2_ with high-glucose DMEM (FUJIFILM Wako Pure Chemical) supplemented with 10% fetal bovine serum (FBS, Sigma), 2mM L-Alanyl-L-Glutamine (FUJIFILM Wako Pure Chemical Corporation), and 100U/mL penicillin-streptomycin (Gibco).

### Induction of cellular senescence

Young cells (population doubling level: PDL 32-46) were used as controls. For replicative senescence, WI-38 cells were passaged until they lost the ability to proliferate and became senescent around PDL 52.

For the plasma membrane damage-dependent senescence (PMDS), DNA damage response-dependent senescence (DDRS), and calcium influx-dependent senescence (CIS), WI-38 cells were treated with 0.0090% SDS, 250 nM doxorubicin (Cayman Chemical), or 75 mM KCl-containing medium, respectively, for 24 hr. The day after treatment, cells were washed twice with fresh medium, then cultured up to 16 days with medium change for every four days. Samples were then collected from the non-treated cells and the treated cells at 0 day (24 hr of treatment), and 1, 2, 3, 4, 5, 6, 8, and 16 days after treatment for PMDS, DDRS, and CIS.

### Protein extraction and Western blotting

Cells were lysed with 2X Laemmli buffer (4% SDS, 20% glycerol, 0.02% bromophenol blue, 0.125M Tris-HCl pH 6.8, 2% 2-mercaptoethanol). The lysate was boiled at 98°C for 3 min and sonicated by BIORUPTOR II (BM equipment) (20 cycles of 20 sec on/30 sec off at high power). The lysate was clarified by centrifugation at 15,000 rpm for 15 min. Protein concentrations were determined by using Qubit protein assay kit (Thermo Fisher Scientific). Five microgram protein samples were loaded onto 4-15% Tris-Glycine gels (Mini-PROTEAN TGX Precast Gels, Bio-Rad) and transferred onto PVDF membranes. The blotted membranes were blocked by Bullet Blocking One (Nacalai) for 5 min and incubated with primary antibodies. After quick rinses with Milli-Q water (5 times) and one 5 min wash with TBS-T, the membranes were incubated with HRP-conjugated secondary antibody for 1 hr at RT. After the same washes as following primary antibody, blots were developed using ECL prime (Amersham) and detected by ChemiDoc MP Imaging System (Bio-Rad). For immunodetection using the same membrane, the antibodies on the membrane were stripped by Restore Western Blot Stripping Buffer (Thermo Fisher Scientific). Antibody conditions were as follows: Anti-p53 (sc-126, Santa Cruz) 1:2000 in Bullet ImmunoReaction Buffer (Nacalai) (1 hr at RT) followed by 1:2000 secondary antibody, Anti-p21 (ab109520, Abcam) 1:5000 in 0.5X Bullet Blocking One in TBS-T (1 hr at RT) followed by 1:5000 secondary antibody, Anti-p16 (ab108349, Abcam) 1:1000 in 0.5X Bullet Blocking One in TBS-T (overnight at 4°C) followed by 1:5000 secondary antibody, Anti-GAPDH (5174, Cell Signaling Technology) 1:5000 in 0.5X Bullet Blocking One in TBS-T (1 hr at RT) followed by 1:10000 secondary antibody. The band intensities were quantified by ImageJ software (NIH) and normalized by the SDS D-1 sample (10th lane) on each membrane (Fig. S1).

### RNA extraction

Total RNA was extracted and purified according to the manufacturer’s instructions using TRIzol (Thermo Fisher Scientific) and RNA Clean and Concentrator-5 kit (Zymo Research). The concentrations of extracted RNA were determined by using a NanoDrop One^C^ spectrophotometer (Thermo Fisher Scientific, Wilmington, DE, USA) and Qubit fluorometer (Thermo Fisher Scientific). The integrity of RNA was checked using an automated electrophoresis system, 4200 Agilent TapeStation (Agilent).

### mRNA sequencing

All samples were sequenced by Novaseq6000. Paired-end cDNA libraries were sequenced with a 2×150-bp configuration. Each sample was sequenced at a depth of more than 20 million paired-end reads.

### Bioinformatics analyses

Identification of SASP components was performed by comparing the significantly expressed genes in each time course of PMDS, CIS, and DDRS with the SASP Atlas (12). SASP Atlas contained 1476 SASP factors expressed from different fibroblast cells induced by different senescent inducers, namely irradiation (IR), inducible RAS overexpression (RAS), and atazanavir treatment (ATV; a protease inhibitor used in HIV treatment) from different fibroblast cells. The overlapping SASP factors in PMDS, CIS, DDRS, and SASP Atlas were shown in a Venn diagram.

### Gene ontology (GO) pathway enrichment analyses

For functional classification, Gene Ontology (GO) pathway enrichment analysis was performed by the Protein Analysis Through Evolutionary Relationships (PANTHER) classification system. PANTHER includes all annotations provided by the Gene Ontology Consortium (http://geneontology.org) to enable identification of genes or gene sets related to the general pathways. It also provides additional information on the genes such as their specific molecular functions, roles in biological processes, and roles as cellular components (13).

### Pathway and biological functions analyses using Ingenuity Pathway Analysis (IPA) software

Ingenuity Pathway Analysis (IPA) (Ingenuity Systems, Redwood City, CA, USA) software (www.ingenuity.com) was used to predict networks that are affected by the differentially expressed genes in PMDS, CIS, and DDRS. Details of the identified genes, their quantitative expression values, and p-values were imported into the IPA software. Firstly, the ‘Core Analysis’ function included in IPA was used to interpret the data in the context of biological processes, pathways, and networks (14).

Both up- and downregulated genes were defined as value parameters for the analysis. Each gene identifier was mapped to its corresponding protein object and was overlaid onto a global molecular network developed from information contained in the Ingenuity Knowledge Base. A network of proteins was then algorithmically generated based on their connectivity. Right-tailed Fischer’s exact test was used to calculate a p-value to indicate the probability that of each biological function being assigned to the network by chance.

Next, the IPA comparison analysis was performed to compare the similarity and difference among the affected pathways and biological functions in the whole time course of PMDS, CIS, and DDRS. The analyses were set up in accordance with our established comparison analyses (15). A heat map of biological functions affected by all stresses was generated to show activity increase (orange bar) and decrease (blue bar) based on the activation z-scores. The upstream regulators were further identified from the significant up- and downregulation of SASP factors during PMDS, CIS, and DDRS and the molecular interactions between the upstream regulators and the downstream SASP factors in the canonical pathways were illustrated. The upstream regulators with activation regulation were shown in orange nodes while those of inhibition regulation were shown in blue nodes. The downstream SASP factors were shown in red (upregulation) and green (downregulation).

### Quantitative reverse-transcription polymerase chain reaction analyses (qPCR)

Complementary DNA (cDNA) was synthesized using a Superscript™ IV VILO Master Mix reagents (Thermo Fisher Scientific) according to the manufacturer’s instructions. The expression of target genes was determined using a QuantStudio 1 real-time PCR system (Thermo Fisher Scientific). PCR amplification was performed using a PowerTrack SYBR Green Master Mix (Thermo Fisher Scientific) with primer pairs as listed in Table S1. PCR was performed in fast cycling mode with the following thermocycling parameters; 2 min of initial DNA polymerase activation and DNA denaturation at 95°C followed by 40 cycles of denaturation at 95°C for 5 sec and primer annealing at 60°C for 30 sec. Melt curves of the real-time PCR products were analysed from 60°C to 95°C. Differences in gene expression, expressed as fold-change, were calculated using the ΔΔCt method, where *Actb* and *Gapdh* were used as reference genes for normalizing the expression.

### Statistical analyses

Results are expressed as means ± standard deviation. The data were statistically analysed using SPSS software, version 21.0 (IBM Corp., Armonk, NY, USA). One-way analysis of variance (ANOVA) and Tukey-Kramer test were used to compare means among groups of different treatments.

NormFinder software was used to determine the stability values of reference genes, *Actb* and *Gapdh*, which were used to normalize gene expression values (16). Venn diagrams were generated using the open access application, Venny (17). The nested scatter plot was used to observe the values for each replicate in the samples used in the experiment (GraphPad Prism).

## RESULTS

### Identification of SASP factors that are differentially expressed in each senescent cell subtype

We previously found that transient plasma membrane damage induced by SDS triggers premature, DNA damage-independent senescence (PMDS) in the normal human fibroblast cells, WI-38 and HCA2 (11). To compare the time-resolved SASP transcriptomic profile in SDS-dependent senescence and other senescence subtypes, we performed mRNA sequencing (mRNA-seq). A successful senescence induction of PMDS and other senescence subtypes were confirmed by the SA-β-gal staining and western blot of senescent factors, p53, p21, and p16, (Fig. 1A-D, S1).

**Figure 1.**
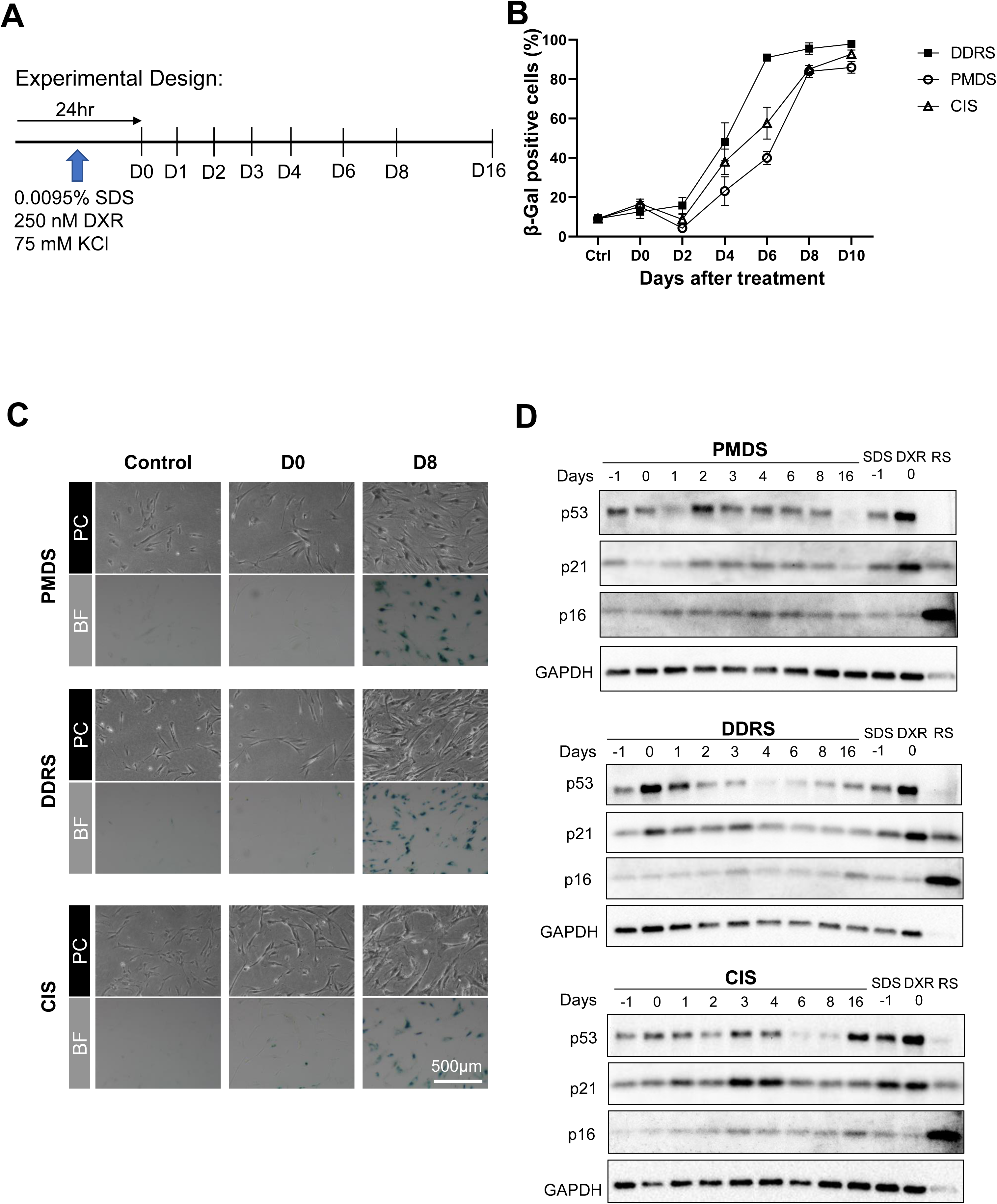
PMD-induced senescence is comparable to other senescence inducers. A. Time phase of senescence inductions in WI-38 cells induced by SDS (PMDS), KCl (CIS), and DXR (DDRS). B-C. Graph and images show the SA-β-gal staining and D. Western blot of senescent factors, p53, p21, and p16, confirming a successful senescence induced by RS, PMDS, CIS, and DDRS.

We performed mRNA-seq using the samples collected at each time point during senescence progression (Fig. 1A). SASP factors were identified using SASP Atlas, which includes the SASP proteins, their upstream regulators, and the proteins in the plasma of aged individuals (12) (Fig. S2A-D). We could successfully identify SASP factors from all differentially expressed genes (in PMDS, 431-772 SASP factors were identified at each time point; CIS, 399-776; DDRS, 445-706; RS, 564). We compared the SASP factors that were differentially expressed in PMDS, CIS, and DDRS at each time point. We found that the SASP factors are more diverse at the early phase (Day 1-3), although the SASP factors are more common at the later phase (Day 8 and 16) (Fig. 2A-B, Fig. S3).

**Figure 2.**
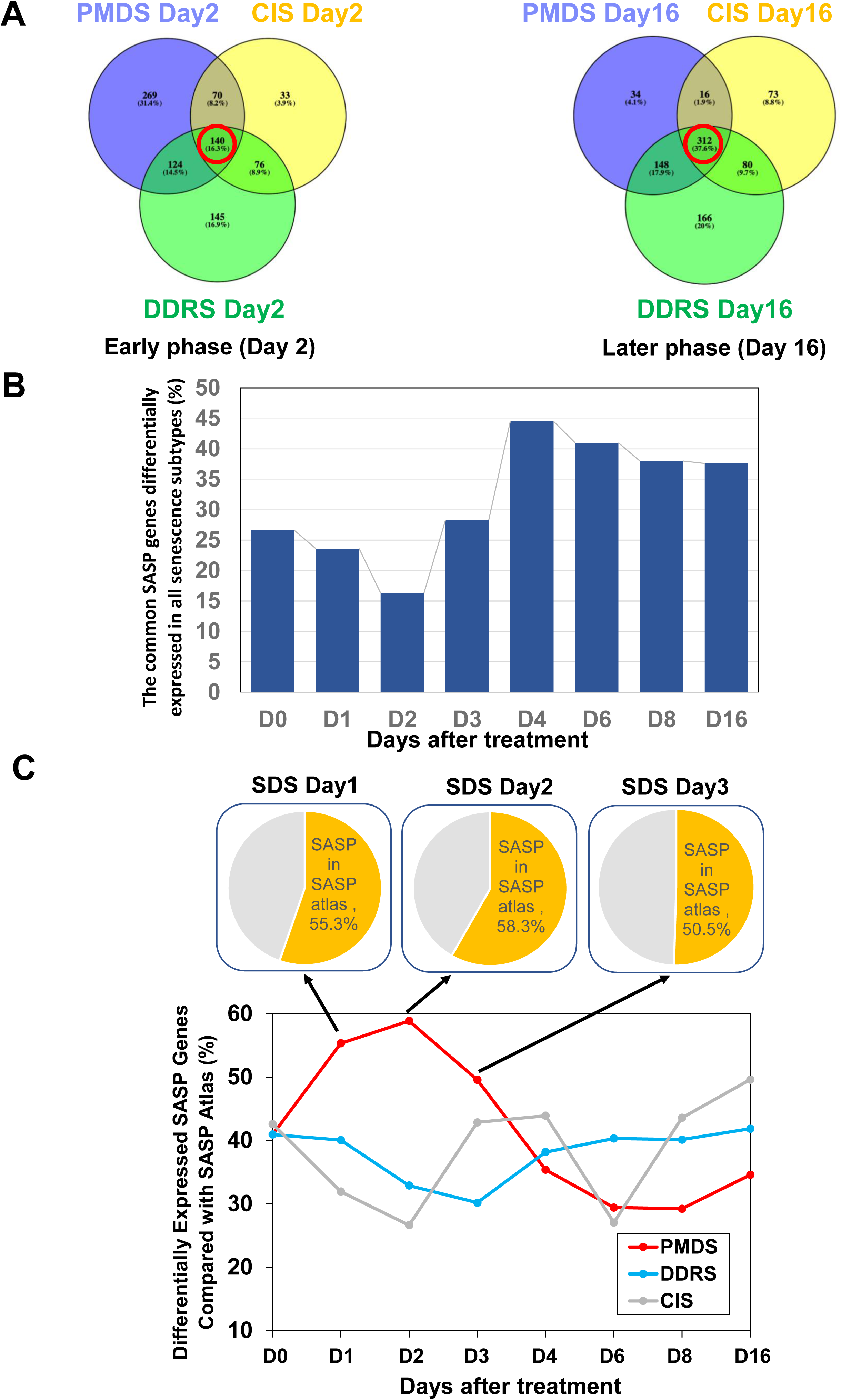
SASP factors are homogenous at the later phase of senescence induction. A. Venn diagram shows the overlapping genes (red circle intersections) in PMDS, CIS, and DDRS on Day 2 and Day 16. B. Graph shows the number of overlapping genes in all senescence inducers increased from the early phase (Day 0) to the later phase (Day 16). C. PMD-dependent SASP factors increased at the early phase. Pie chart and graph show a striking increase of SASP factors at the early phase of PMDS (Day 1, 2, 3) but not in CIS and DDRS.

The number of SASP factors differentially expressed in PMDS strikingly increased during Day 1-3 (Fig. 2C, red). In CIS, the number of SASP factors fluctuates (Fig. 2C, grey). In contrast, relatively constant numbers of SASP factors were identified in DDRS (Fig. 2C, blue). To check if the SASP-associated pathways are enriched in the early phase of PMDS, PANTHER classification system was used. At the early phase (Day 2), when PMDS cells expressed the highest number of SASP factors (Fig. 2C), GO enrichment analysis showed a significant enrichment of the SASP-associated pathway, namely “Inflammation mediated by chemokine and cytokine signaling” (Fig. 3A). However, this pathway was not listed within the top five mostly enriched pathways in CIS and DDRS on Day 2, consistent with the lower number of SASP factors identified in these conditions (Fig. 2C). Moreover, consistent with the increase of SASP factors at Day 16, the pathway was also enriched in CIS at Day 16 (Fig. 3B). This pathway was enriched in the early phase of PMDS (Day 2 and 3) and CIS (Day 3) (Fig. 3A, S4) and the later phase of CIS (Fig. 3B). These results support the interpretation that SASP is upregulated at early PMDS.

**Figure 3.**
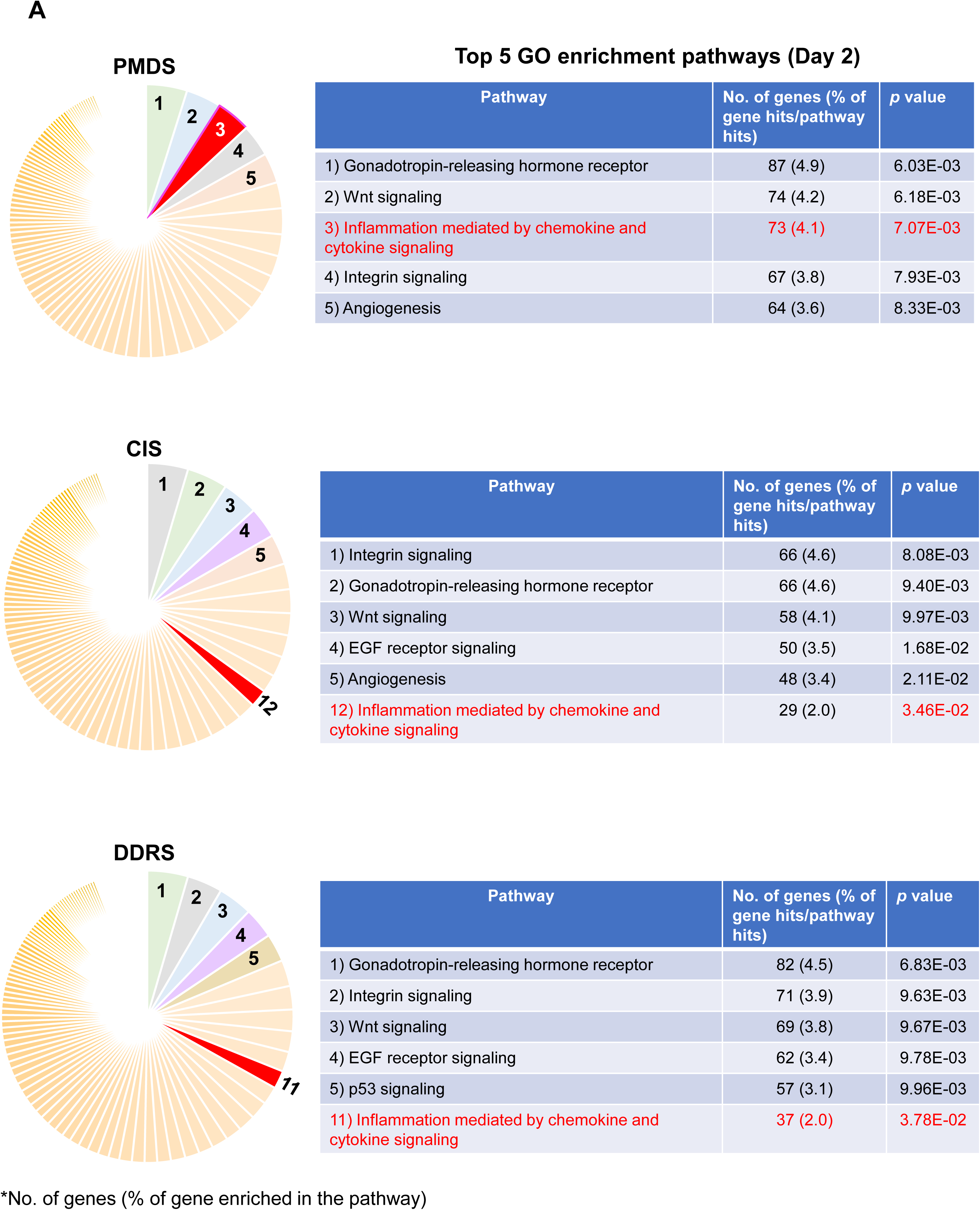
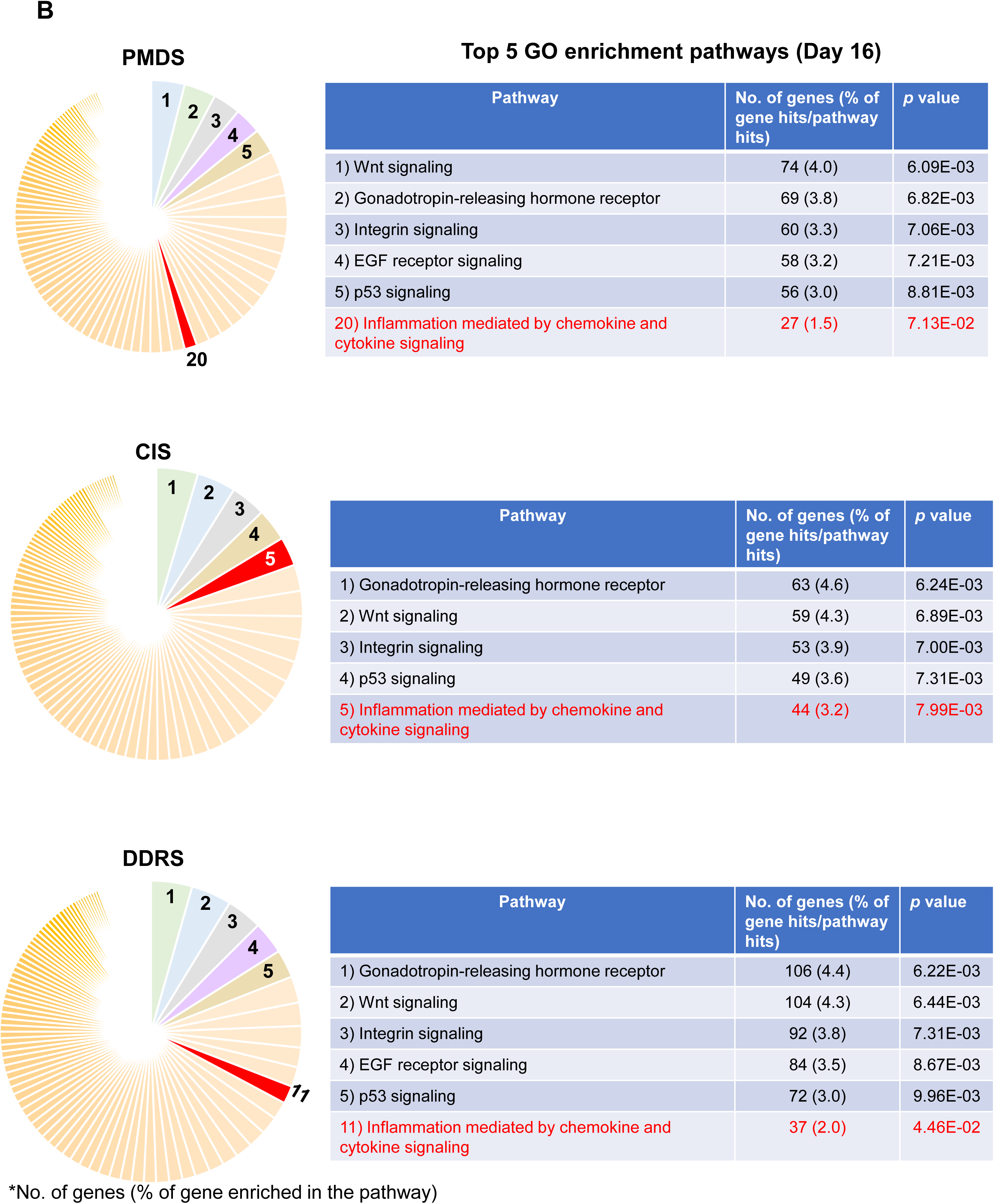
SASP-associated pathway was enriched only at the early phase of PMDS and the later phase of CIS. Pie chart and tables show the top five hit pathways for the enrichment of Gene Ontology (GO) SASP-associated pathway only in the early phase of PMDS (Day 2) (A) and the later phase of CIS (Day 16) (B). The GO was analysed using PANTHER classification system (http://geneontology.org).

To test if the enrichment of the SASP-related “Inflammation mediated by chemokine and cytokine signaling” pathway is relevant, we further identified the sub-categories of its downstream targets. Day 3 data was subjected to the analysis because the SASP expression was relatively high both in PMDS and CIS on the day (Fig. 2C). The downstream categories, extracellular matrix (ECM) protein, NF-κB regulator, calcium regulator, phospholipase C, and cell cycle, were all enriched on Day 3 in PMDS and CIS (Fig. 4A), consistent with the increase of SASP factors at Day 3 in PMDS and CIS (Fig. 2C). Further, the downstream targets of ECM protein, NF-κB regulator, calcium regulator, and phospholipase C were also enriched at the early phase of PMDS and CIS (Day 1-2) (Fig. S5A-B).

**Figure 4.**
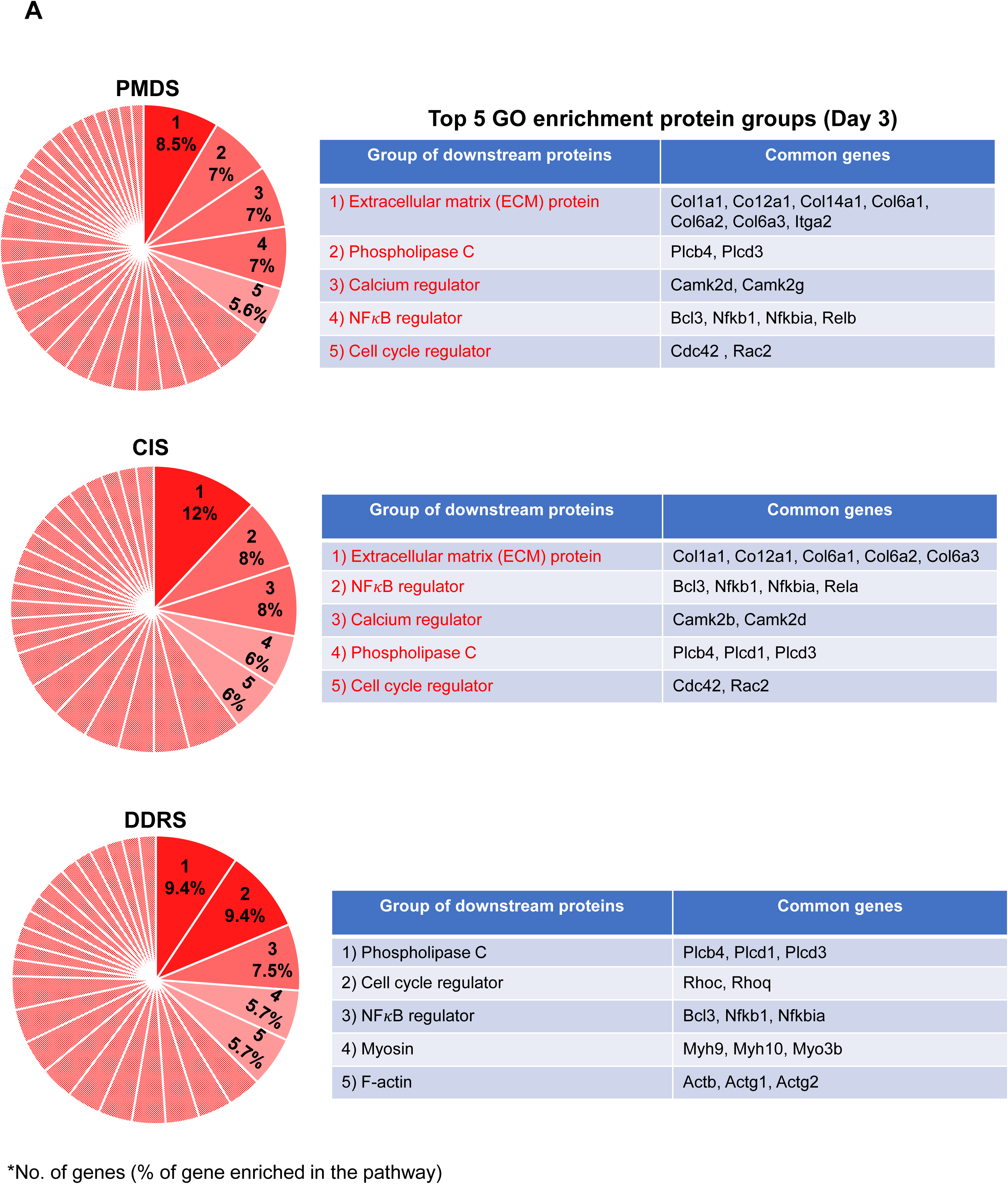
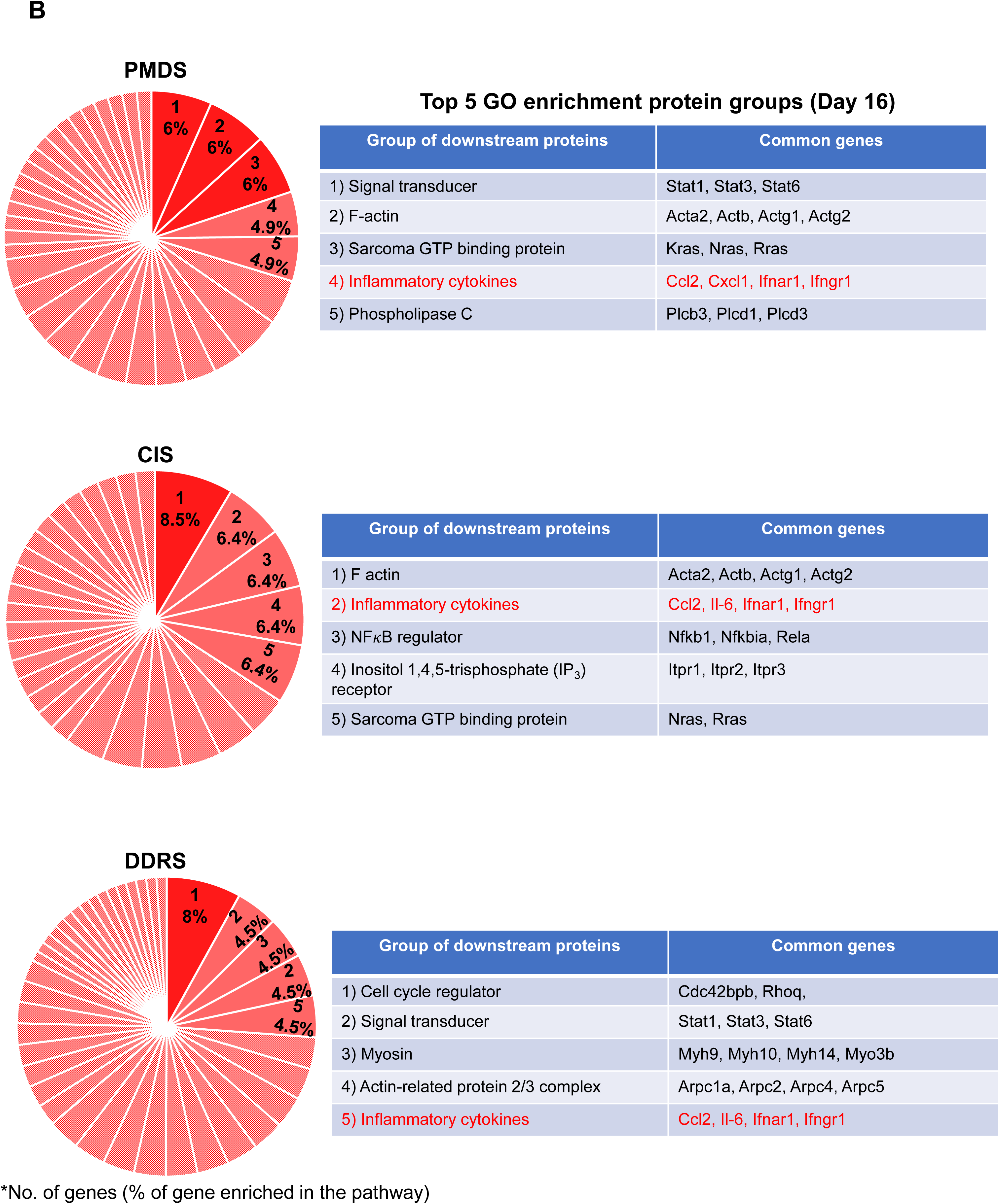
Similar downstream targets were identified in the SASP-associated pathways at the early phase of PMDS and CIS. Pie chart and tables show the top five hit categories for the enrichment of Gene Ontology (GO) for the sub-categories of downstream targets under the SASP-associated pathway at the early phase (Day 3) (A) and the later phase (Day 16) (B) of PMDS, CIS, and DDRS. The GO was analysed using PANTHER classification system (http://geneontology.org).

These results strongly suggest that the SASP-related pathways are differentially regulated at the early phase of PMDS. In contrast, at the later phase (Day 16), the downstream pathways were uniformly enriched in all senescent cell subtypes (Fig. 4B), which might reflect the homogeneous pattern of SASP expression profiles at the later phase of senescence. These results suggest that while different senescent cell subtypes express diverse SASP factors at the early phase, all senescent cell subtypes express relatively uniform SASP factors at the later phase (Fig. 2, S3).

### Top canonical pathways affected by the significant gene expression in PMDS

To identify the pathways activated/inhibited in each senescent cell subtypes, we performed the pathway comparison analyses using IPA software. The benefit of using this software is that we can use cell type-specific databases, such as the one for normal human fibroblasts. We also identify the top canonical pathways associated with a dataset of significantly expressed genes in PMDS, DDRS, CIS, and RS, which uses the Fisher’s exact test to ascertain pathway enrichment. We first focused on the top canonical pathways in PMDS and compared them with CIS and DDRS. The pathway comparison analysis identified 21 top canonical pathways affected in PMDS from Day 0 to 16 whose activation z-score values are greater than 2 (activation) or smaller than -2 (inhibition) (Fig. 5). The IPA’s z-score indicates a predicted activation or inhibition of a pathway/gene, where a positive z value connotates an overall pathway’s activation and a negative z value connotates an overall pathway’s inhibition.

**Figure 5.**
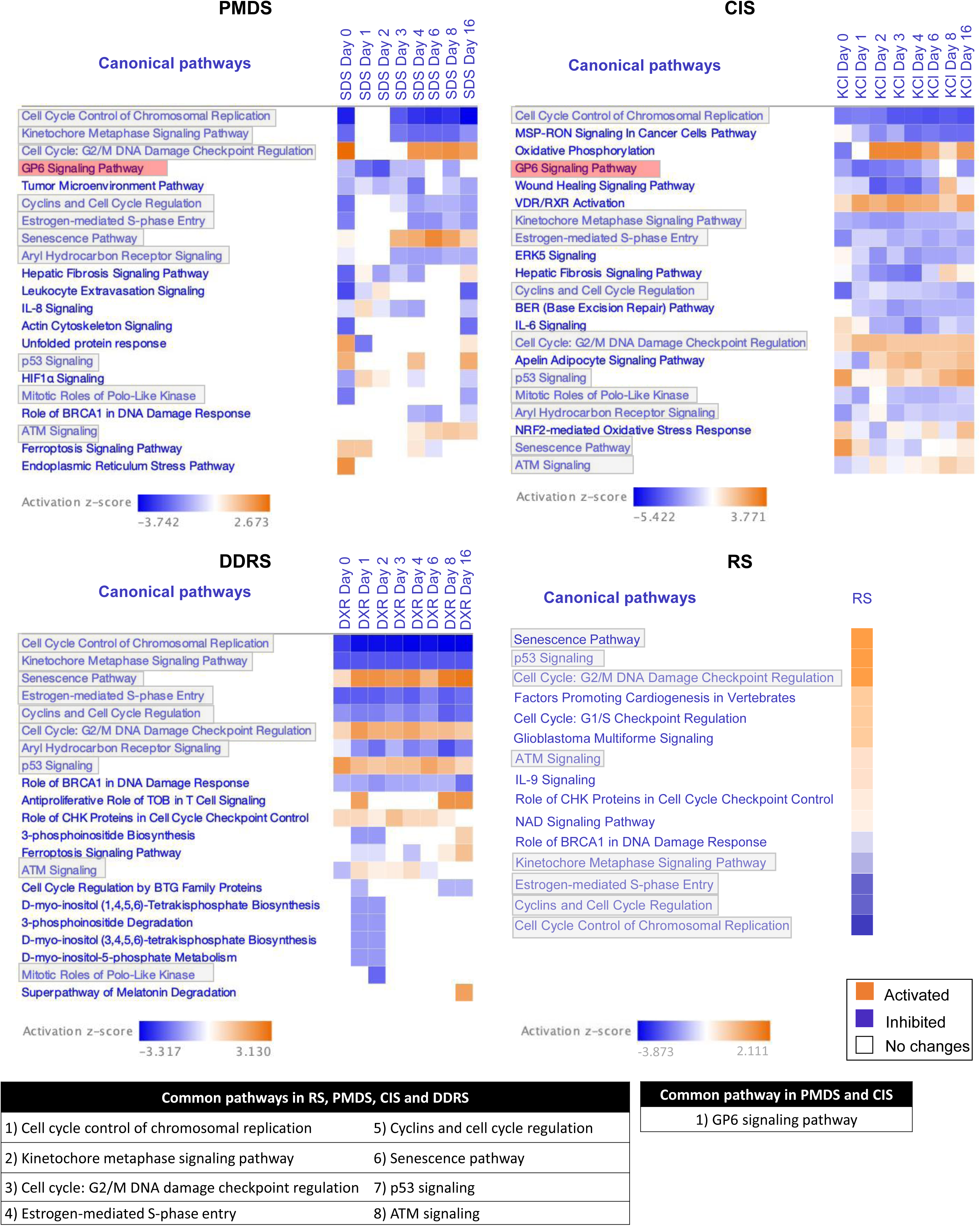
Top canonical pathways are generated from the significant gene expression in response to PMDS, CIS, DDRS, and RS. Heat maps generated from IPA comparison analysis show top pathways affected by the gene expression in each time phase in PMDS, CIS, DDRS, and RS. Orange and blue bars indicate positive and negative activation z-score respectively, while white bar indicates no activation.

According to the same cut-off z-score values, we also identified 21 top canonical pathways in CIS and DDRS. Ten out of 21 pathways were common among PMDS, CIS, and DDRS (Fig. 5, gray highlights, and the bottom left table). The common features of PMDS, CIS, and DDRS are inhibition of cell cycle-related pathways and activation of the cell senescence-related pathways. In a positive control, replicative senescence (RS), these two features were identified as well, suggesting that the core senescence mechanisms were common in all senescent cell subtypes (Fig. 5).

Strikingly, we found that the central senescence mechanisms, namely senescence pathway and the p53 pathway, are regulated differently in different senescent cell subtypes. The activation/inhibition of these pathways are comparable to that of other senescent cell subtypes only after Day 4. These results suggest that the progression of senescence is slower in PMDS than DDRS. Indeed, this interpretation is consistent with SA-β-Gal staining (Fig. 1B). In PMDS, the senescence pathway was activated only at Day 3 and the subsequent days. In CIS, the senescence pathway was activated at Day 0, 1, and 16. In contrast, in DDRS, the senescence pathway was constantly activated. We also found that, in PMDS, p53 signaling was only activated at Day 0, 4, and 16. In CIS, p53 signaling was strongly activated only in Day 0, 3, 8, and 16. In contrast, in DDRS, p53 signaling was activated during all phases (Fig. 5). These results suggest that the senescence mechanism mediated by the p53 pathway is diverse in PMDS, CIS, and DDRS, although all treatments ultimately induce senescence (Fig. 1).

### GPVI signaling pathway was inhibited via SASP in PMDS and CIS but not in DDRS

To identify the unique mechanism underlying PMDS, we sought the pathways that are specifically altered in PMDS. The IPA pathway comparison analyses identified “GPVI signaling pathway”. This pathway was strongly inhibited in the early phase of PMDS and CIS (Fig. 5, red highlights).

Upregulation of SASP factors and enrichment of SASP-related pathways at the early phase of PMDS led us to hypothesize that SASP factors might suppress “GPVI signaling pathway”. To test this idea, we performed the pathway comparison analyses using the list of SASP factors identified in PMDS, CIS, and DDRS (Table S2).

Using our SASP hit list, we found that seven common canonical pathways were activated in PMDS, CIS, and DDRS, namely wound healing signaling pathway, RHOGDI signaling, signaling by Rho family GTPases, integrin signaling, regulation of actin-based motility by Rho, leukocyte extravasation signaling, and HMGB1 signaling (Fig. 6, right table).

**Figure 6.**
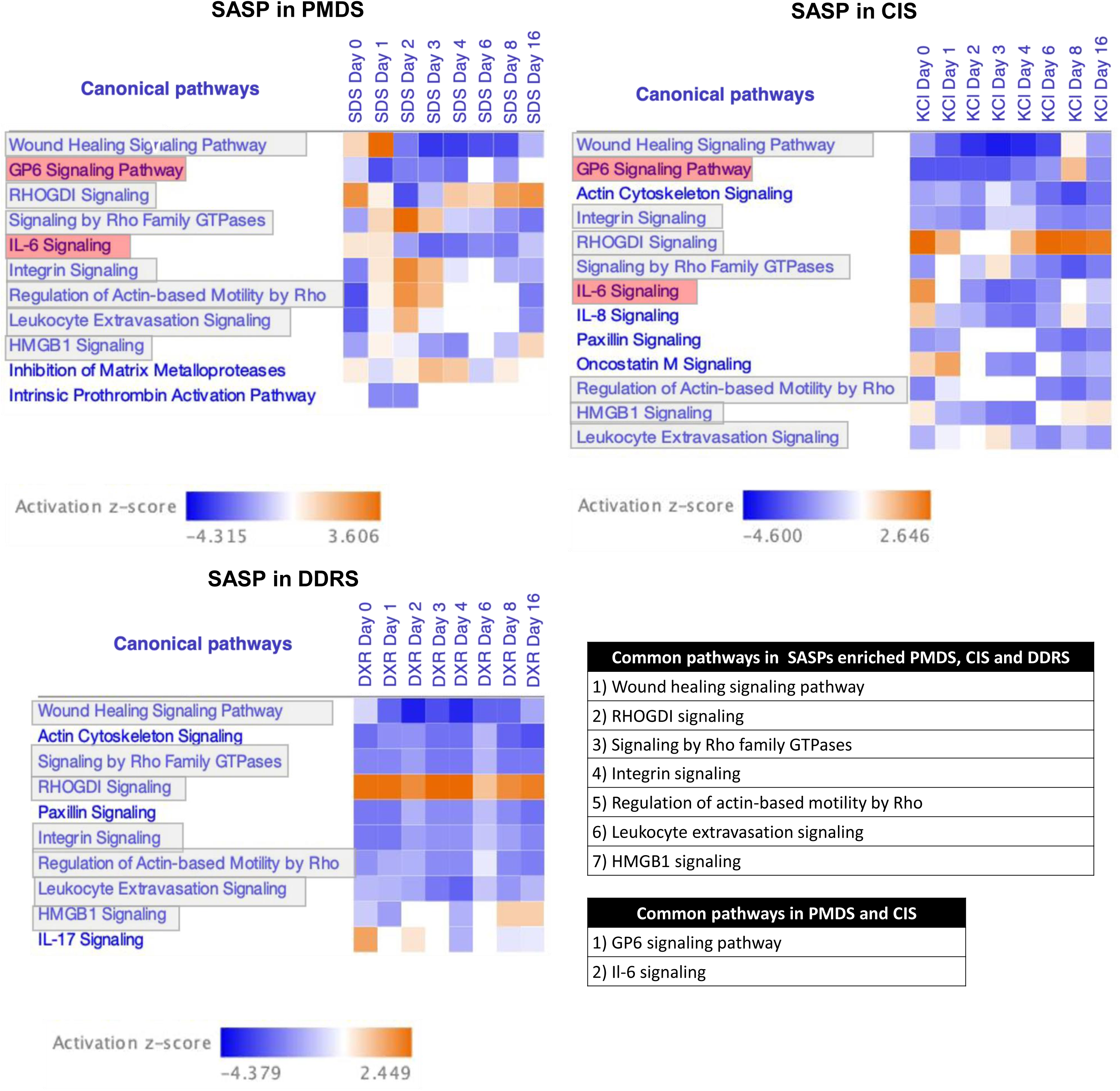
Top canonical pathways are generated from the significant SASP factors expression in response to PMDS, CIS, DDRS, and RS. Heat maps generated from IPA comparison analysis show top pathways affected by the SASP gene expression in each time phase in PMDS, CIS, DDRS, and RS. Orange and blue bars indicate positive and negative activation z-score respectively, while white bar indicates no activation.

We found that the GPVI signaling pathway was inhibited strongly at the early phase of PMDS, both in the SASP factor hit lists and the entire differentially expressed gene hit lists. (Fig. 5 and 6). Similar results were obtained at the early phase of CIS (Fig. 5 and 6). In addition, Il-6 signaling and RHOGDI signaling were analogously regulated at the early phase of PMDS and CIS. However, the major difference exists in the wound healing signaling pathway, which was induced only at the early phase of PMDS. In contrast, Oncostatin M signaling was activated exclusively at the early phase of CIS. These results suggest that the inhibition of GPVI signaling pathway is a common feature of early PMDS and CIS, but it could potentially be regulated via different SASP factors.

We also observed that RHOGDI and HMGB1 signalings were commonly activated at the later phase of PMDS, CIS, and DDRS in the SASP factors hit list (Fig. 6). These results suggest that the upregulation of common SASP factors at the later phase might be the underlying mechanism for these pathways activation.

By using the IPA’s upstream regulator analysis, we next explored the potential upstream pathways associated with the inhibition of the GPVI signaling pathway at the early phase of PMDS. At the early phase of PMDS, the activation of wound healing signaling pathway showed the most significant association with the inhibition of GPVI signaling pathway (p < 1.1 x 10^-9^) (Fig. 7, middle right), followed by Il-6 signaling (p < 4.2 x 10^-7^) (Fig. 7, upper left). Activation of wound healing signaling pathway was specifically associated with the activation of RHOGDI signaling. IPA predicted that RHOGDI signaling indirectly inhibits GPVI signaling pathway via its strong activation on the wound healing signaling pathway (Fig. 7, lower right). In these GPVI signaling-associated pathways, the SASP factors are the major (>50%) downstream targets in all pathways except for one (integrin signaling) (Fig. 7, upper right, SASP:31%). In addition, overlapped SASP factors, namely *Cxcl8*, *Il-1α*, *Il-1β*, *Il-6*, *Mmp1,* and *Mmp3*, were the downstream targets of four out of seven pathways (inhibition of matrix metalloproteinases, leukocyte extravasation signaling, wound healing signaling pathway, and Il-6 signaling) (Fig. 7, gene names in red). These results are consistent with our earlier conclusion that the SASP factors contribute to the inhibition of the GPVI signaling pathway.

**Figure 7.**
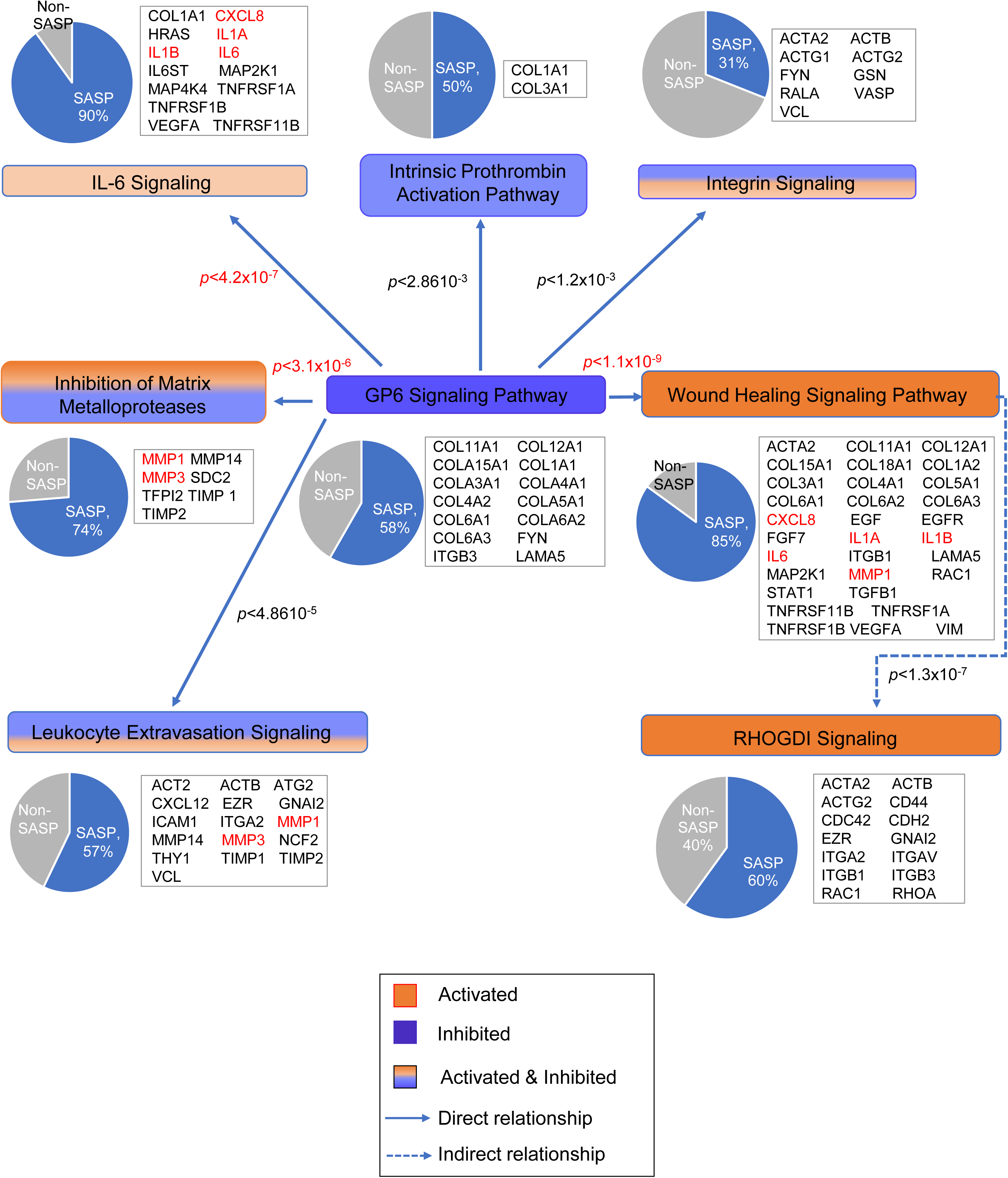
GPVI signaling pathways are implicated in various SASP factors regulated at the early phase of PMDS. Diagram shows the significant GPVI signaling-associated pathways affected by PMDS analysed by IPA software. Pie chart shows that SASP factors are major downstream targets in these pathways. Common SASP factors, namely *Il-6*, *Mmp1*, and *Mmp3* (red fonts), are the downstream genes in most of the GPVI signaling-associated pathways. Orange and blue bars indicate positive and negative activation z-score respectively.

Analogous to PMDS, the inhibition of GPVI signaling at the early phase of CIS was significantly associated with the wound healing signaling pathway, Il-6 signaling, leukocyte extravasation signaling, and integrin signaling (Fig. S6). However, unlike PMDS, the activation of Il-6 signaling and Il-8 signaling showed higher significant association with the inhibition of GPVI signaling pathway (p < 3.7 x 10^-6^ and p < 3.2 x 10^-5^, respectively, Fig S6 upper left and upper right). IPA predicted that RHOGDI signaling indirectly inhibits GPVI signaling pathway by activating the Il-8 signaling pathway (Fig. S6, lower right). In addition, Oncostatin M signaling indirectly inhibits GPVI signaling pathway by activating Il-6 signaling (Fig. S6, lower center). Rather than direct regulation by the wound healing signaling pathway as in PMDS, GPVI signaling inhibition in CIS is indirectly regulated by RHOGDI signaling and Oncostatin M signaling. Common SASP factors were also implicated in the GPVI signaling-associated pathways during the early phase of CIS, namely *Cxcl8*, *Il-6*, *Mmp1,* and *Mmp3* (Fig. S6, gene names in red). These results suggest that the inhibition of GPVI signaling pathway in PMDS and CIS involves common SASP factors, but the molecular mechanisms are partly different.

### Transient upregulation of wound healing SASP factors, *Il-1α*, *Il-1β*, *Il-6*, *Cxcl8*, *Mmp1,* and *Mmp3*, could be the upstream regulator of the GPVI signaling pathway at the early phase of PMDS

In PMDS, the wound healing signaling pathway was the most significantly activated on Day 1 (Fig. 6). Several pathways were the most significantly activated on Day 2 (signaling by Rho family GTPases, integrin signaling, regulation of actin-based motility by Rho, and leukocyte extravasation signaling) (Fig. 6). RHOGDI signaling was activated on Day 0 and Day 4-16. These results raise a possibility that at the early phase of PMDS, the cells may play a specific role in vivo via these SASP-related pathways. We speculated that the different SASP factors contribute to the different pathways regulation at the early phase of PMDS.

Therefore, we aimed to identify the upstream regulators of SASP factors at the early phase of PMDS (Fig. 8A). This analysis could potentially suggest the mechanism underlying PMDS-specific functions in vivo. Briefly, we can identify the cascade of upstream transcriptional regulators that can explain the observed gene expression changes in the early phase of PMDS, which can help illuminate the biological activities occurring in the PMDS-induced cells. The molecular interactions between the upstream regulators and the other SASP factors is displayed graphically as nodes (genes) and lines (the biological activation or inhibition relationship between the genes). The activated upstream regulators were shown in orange nodes while the inhibited ones were shown in blue nodes. The downstream SASP factors were shown in red nodes (upregulation) and green nodes (downregulation).

**Figure 8.**
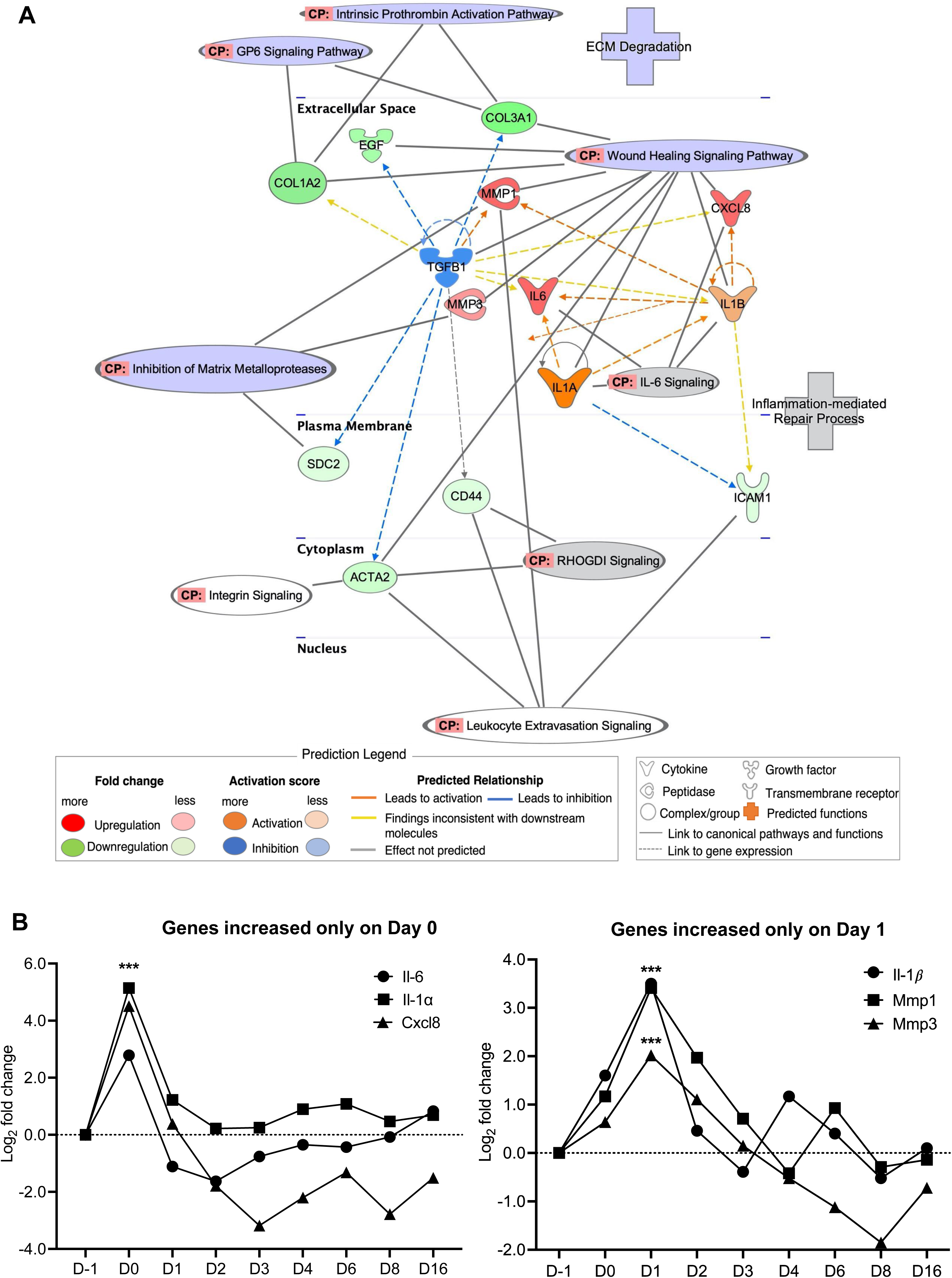
GPVI signaling is inhibited by transient increase of wound healing SASP factors early after PMDS. IPA graphical representation shows the upstream regulators of GPVI signaling inhibition mainly involved in wound healing SASP factors (A). The molecular interactions between the upstream regulators and the other SASP factors are displayed graphically as nodes (genes) and dotted lines (the biological activation or inhibition between the genes). Nodes in red indicate upregulated genes while those in green represent downregulated genes. Graph shows the expression of wound healing SASP factors regulated in PMDS from Day 0-16. *Il-1α*, *Il-6*, and *Cxcl8* were transiently upregulated at Day 0, while *Il-1β*, *Mmp1*, and *Mmp*3 were upregulated at Day 1 (B).

The analyses predicted that the upregulation of *Il-1α* and *Il-1*β (orange nodes) and the downregulation of *Tgfb1* (blue node) were the upstream regulators of the SASP factors at the early phase of PMDS (Fig. 8A). The molecular interactions showed that the upregulation of *Il-1α* and *Il-1*β with the downregulation of *Tgfb1* further activated the SASP factors, *Il-6*, *Cxcl8*, *Mmp1,* and *Mmp3*, that are implicated in wound healing signaling (Fig. 8A, upper right, purple) and Il-6 signaling (Fig. 8A, middle right, grey) pathways. Those SASP factors prominently downregulated the SASP factors involved in the GPVI signaling pathway (Fig. 8A, upper left, purple). The pathways in this network are closely associated with the ECM degradation and inflammation-mediated repair process.

These results were confirmed in our mRNA-seq data sets, where we observed transient upregulations of *Il-1α, Il-1β*, *Il-6*, *Cxcl8*, *Mmp1,* and *Mmp3* during the early phase of PMDS (Fig. 8B).

We also observed the transient upregulation of common SASP factors, namely *Il-6*, *Cxcl8*, *Mmp1,* and *Mmp3*, at the early phase of CIS (Fig. S7B). Those SASP factors were prominently implicated in Il-6, Il-8, and Oncostatin M signaling, which indirectly associated with wound healing signaling pathway (Fig. S7A and Fig. S6). These results suggest the inhibition of GPVI signaling at the early phase of PMDS and CIS was regulated by the transient upregulation of common SASP factors.

### Persistent upregulation of *Ccl2* and *Il-6* are the upstream regulators of cell senescence at the later phase of PMDS, CIS, and DDRS

To identify the senescence subtype-specific regulatory network, we compared the upstream regulators at the later phase of PMDS, CIS, and DDRS. As shown in the molecular interactions, *Ccl2* is the main upstream regulator (Fig. 9A, orange node) that targets downstream SASP factors in HMGB1 signaling, wound healing signaling pathway, leukocyte extravasation signaling, and inhibition of matrix metalloproteases. In contrast to the transient expression of wound healing SASP at the early phase, we observed a persistent upregulation of *Ccl2* during the later phase of PMDS at Day 6 and 16 (Fig. 9B).

**Figure 9.**
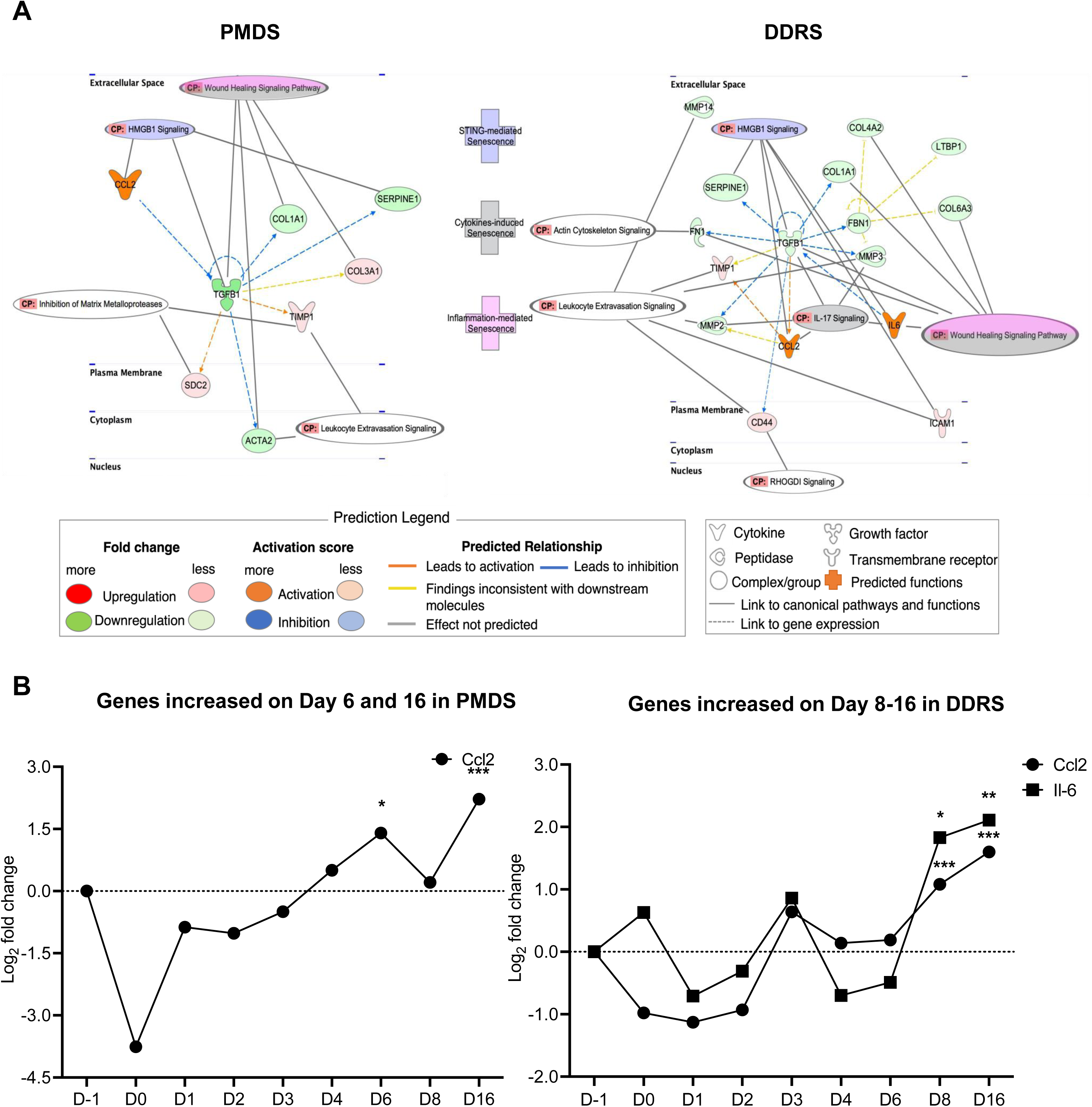
*Ccl2* and *Il-6* induce senescence late after PMDS and DDRS. IPA graphical representation shows *Ccl2* and *Il-6* as the upstream regulators in the later phase of PMDS and DDRS, which induce senescence, particularly by affecting HMGB1 and wound healing signaling pathways (A). The molecular interactions between the upstream regulators and the other SASP factors are displayed graphically as nodes (genes) and dotted lines (the biological activation or inhibition between the genes). Nodes in red indicate upregulated genes while those in green represent downregulated genes. Graph shows the expression of SASP factors, *Ccl2* and *Il-6*, regulated in PMDS and DDRS from Day 0-16. *Ccl2* and *Il-6* were persistently upregulated from Day 6-16 in PMDS and Day 8-16 in DDRS (B).

In the later phase of DDRS, the upregulation of *Il-6* and *Ccl2* (orange nodes) in turn regulated SASP factors. The molecular interactions showed that the upregulation of *Il-6* and *Ccl2* significantly downregulated *Tgfb*1 which eventually led to the regulation of the main downstream SASP factors involved in actin cytoskeleton signaling, wound healing signaling pathway, leukocyte extravasation signaling, Il-17 signaling, and HMGB1 signaling. On the other hand, the downregulation of *Tgfb1* led to the upregulation of the downstream SASP factors in RHODGI signaling (Fig. 9A).

We also observed a persistent upregulation of *Il-6* and *Ccl*2 during the later phase of DDRS after 8 to 16 days (Fig. 9B). This led to activation of HMGB1 signaling which activate during the later phase of DDRS (Fig. 9A).

In the later phase of PMDS and DDRS, *Il-6* and *Ccl2* particularly affected HMGB1 signaling and other pathways that are associated with STING-mediated senescence, cytokines-induced senescence, and inflammation-mediated senescence (Fig. 9A).

Similarly, we observed the persistent upregulation of common SASP factors, *Ccl2* and *Il-*6, later after CIS. In the later phase of CIS, the upregulation of *Tnf* and *Ccl*2 (Fig. S8A, orange nodes), and the downregulation of *Cdkn2a* (Fig. S8A, blue node) were the upstream regulators of SASP factors. The molecular interactions showed that the upregulation of *Il-6* and *Ccl2* significantly affected HMGB1 signaling, wound healing signaling pathway, and leukocyte extravasation signaling that are associated with STING-mediated senescence, cytokines-induced senescence, and inflammation-mediated senescence (Fig. S8A). In addition to the persistent upregulation of *Il-6* (Fig. S7B), we also observed a persistent upregulation of *Ccl2* during the later phase of CIS (8 to 16 days) (Fig. S8B).

Hence, we speculated that, in PMDS, autocrine signaling by SASP factors transiently upregulates pro-inflammatory, wound healing SASP factors, namely *Il-1α, Il-1β*, *Il-6*, *Cxcl8*, *Mmp1,* and *Mmp3*, to activate inflammation-mediated repair/injury responses. In contrast, in DDRS, non-SASP factors might regulate the senescence at the early phase in addition to the SASP persistent upregulation at the later phase.

We also found a transient upregulation of *Stc1*, a newly identified SASP factor (12), early after PMDS and CIS. Stc1 was reported to modulate calcium regulation across cell membranes (18). Hence, we speculated that the transient upregulation of wound healing SASP factors is at least partially induced by Ca^2+^ influx which is supported by the observation of common wound healing SASP upregulation early after PMDS and CIS (Fig. 8 and S7). On the other hand, the persistent upregulation of *Ccl2* and *Il-6* in the later phase of PMDS, CIS, and DDRS might induce cellular senescence in a paracrine manner by activation of HMGB1 signaling and inhibition of the wound signaling pathway and leukocyte extravasation signaling

### Verification of mRNA-seq results using qPCR analyses

To verify our mRNA-seq results, we performed qPCR. We confirmed that *Actb* and *Gapd*h are the most stable housekeeping genes with the lowest M value (Fig. S9). Therefore, all targeted gene expressions were normalized to the expression of both *Actb* and *Gapdh*.

Our qPCR analyses confirmed the transient upregulation of *Il-6*, *Mmp1*, and *Mmp3* at the early phase of PMDS and CIS (Fig. 10 and S10). In addition to that, the persistent upregulation of *Il-6* and *Ccl2* was also observed in the later phase of PMDS, CIS, and DDRS (Fig. 10 and S10). These findings supported the similar transient and persistent upregulation seen in the mRNA-seq results.

**Figure 10.**
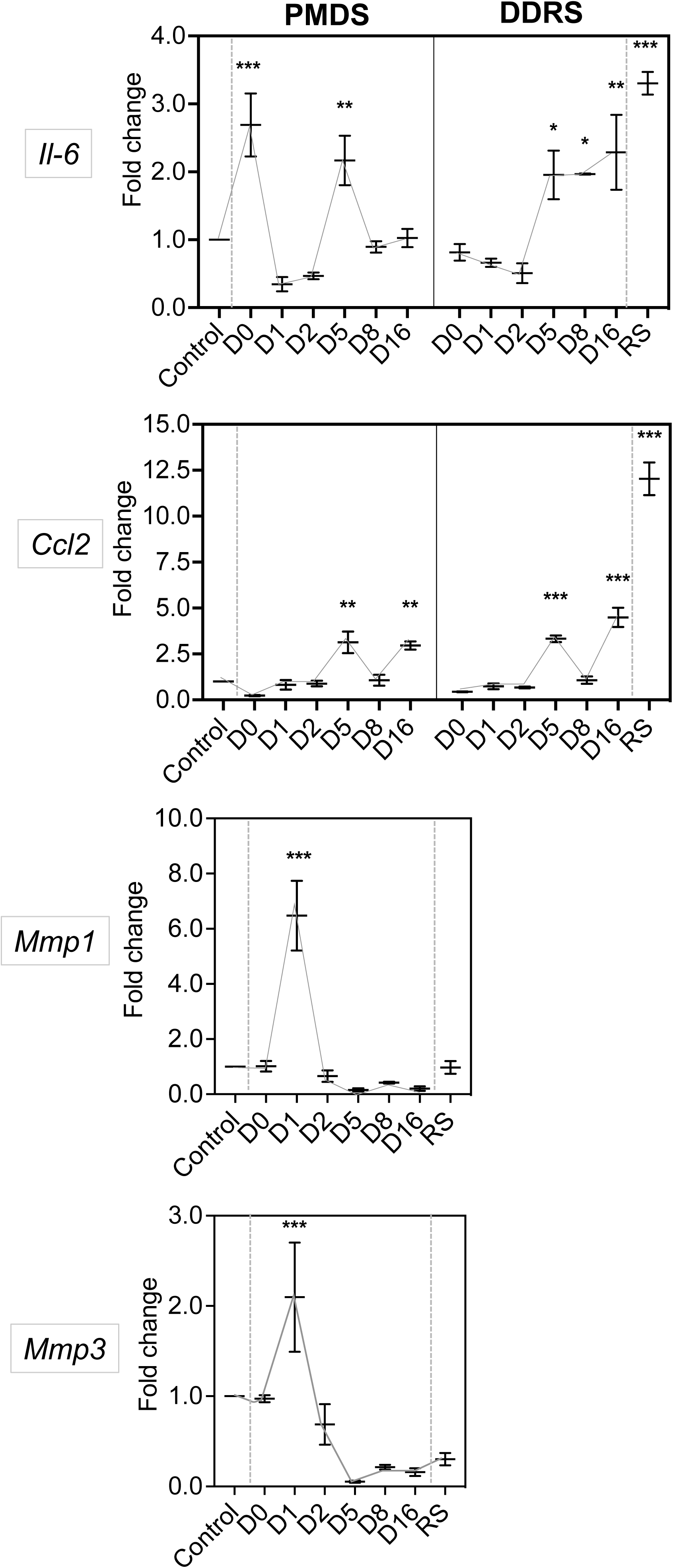
PMDS and DDRS SASP factors upregulation is confirmed by qPCR. Graph shows the mRNA expression of the SASP factors, *Il-6*, *Mmp1*, and *Mmp3*, which were transiently upregulated at Day 0 in PMDS. Graph also shows the mRNA expression of SASP factors, *Ccl2* and *Il-6*, which were persistently upregulated from Day 5-16 in PMDS and DDRS. The expression of all SASP factors was normalized to that of *Actb* and *Gapdh* and calculated as fold changes. Error bars indicate standard deviation. *p<0.05, **p<0.01, ***p<0.001, compared to control, (n=3).

Our qPCR also confirmed the transient upregulation of *Stc1* exclusively at the early phase of PMDS and CIS (Fig. S11). As Stc1 modulates calcium regulation, we speculated that Ca^2+^ influx partially induced the transient upregulation of similar SASP factors in PMDS and CIS. However, further study is needed to reveal the mechanism underlying *Stc1* upregulation early after PMDS and CIS.

## DISCUSSION

In this study, we comprehensively revealed the time-resolved transcriptomic profiles in PMDS, CIS, and DDRS. To our knowledge, our study is the first to identify the SASP factors in the different time courses after PMDS, CIS, and DDRS. We generated the SASP list based on our own mRNA-seq results using multiple senescent cell subtypes (PMDS, CIS, DDRS, and RS) and performed multiple bioinformatics analyses to elucidate the regulatory mechanisms of SASP factors in all senescence phases. A detailed list of SASP factor expressions at each time point with their gene ID, log2 fold change, and *p-*value can be found in the supplementary materials (Table S2). Our analyses identified wound healing SASP factors, *Il-6*, *Mmp1*, and *Mmp3*, that potentially regulate the inflammation-mediated injury/repair response early after PMDS. We also revealed that, at the later phase, PMDS, DDRS, and CIS show persistent upregulation of common SASP factors, *Il-6* and *Ccl2*. Our results suggest that the early stage of senescence is diverse, and the cellular status gradually converges to a relatively uniform pattern.

We found that, in PMDS, most of the upregulated/downregulated pathways returned to the neutral status (comparable to the before the treatment sample) on Day 1 and 2 (Fig. 6). This “gap” period delayed the progression of senescence in PMDS; the senescence pathway was activated only on Day 3 and after. Consistent with this finding, SA-β-Gal positive cells increased slowly in PMDS compared with DDRS; the time point when SA-β-Gal positive cells exceed 50% was Day 6 in DDRS and Day 8 in PMDS (Fig. 1B). These results suggest that the PMDS-specific cellular processes may occur on Day 0-3.

In this work, we utilized the proteomic atlas of aging markers, SASP Atlas (12), to identify the SASP factors in our mRNA-seq data sets. We found a striking increase of SASP factors in the early phase of PMDS (Day 1-3) but not in CIS and DDRS (Fig 2). This finding was further supported by the enrichment of the SASP-associated pathway only at the early phase of PMDS (Fig ). Moreover, we found that the downstream targets in the SASP-associated pathways at the early phase were common between PMDS and CIS. Those downstream targets include ECM protein, NFκB regulator, and calcium regulator. These findings raise a possibility that at least part of the mechanism underlying SASP expression changes in PMDS could be attributed to Ca^2+^ influx. This possibility is consistent with our previous finding that Ca^2+^ influx mediates the upregulation of the p53-p21 axis in PMDS (11).

The GO enrichment analysis indicates that the number of SASP factors differentially expressed in PMDS strikingly increased during Day 1-3 (the early phase) (Fig. 3A). The downstream targets of these SASP factors are overlapping in PMDS and CIS at the early phase (Fig. 4A and Fig. S5), consistent with our idea that Ca^2+^ influx could be one of the mechanisms underlying PMDS.

Intriguingly, we found that the transient upregulation of wound healing SASP factors, particularly *Il-6*, *Cxcl8*, *Mmp1*, and *Mmp3,* are likely to be implicated in the inhibition of the GPVI signaling pathway at the early phase of PMDS (Fig. 7). Moreover, these SASP factors were also implicated in the GPVI signaling pathway in the early phase of CIS (Fig. S6). These results suggest that these SASP factors are important regulators inhibiting the GPVI signaling pathway.

GPVI is a collagen receptor abundantly expressed in platelets. GPVI activates a downstream signaling cascade promoting platelet aggregation and thrombus formation (19). The role of this signaling pathway in plasma membrane-damaged fibroblasts has yet to be explored. However, previous studies linked the GPVI pathway and fibrosis (20). In an in vivo study, the deficiency of platelet GPVI facilitates cutaneous wound repair through a local and temporal vascular leakage leading to increased fibrinogen deposition and reduced leukocyte infiltration. Thus, impaired vascular integrity due to the loss of GPVI is beneficial to wound repair (21). Our results showed the upregulation of proinflammatory cytokines, *Il-l*α, *Il-1*β, *Il-6*, *Cxcl8*, and matrix degrading genes, *Mmp1* and *Mmp3*, that play important roles in wound repair and collagen degradation (22, 23). These results suggest that the inhibition of GPVI signaling was potentially regulated by SASP factors to activate wound repair in vivo. However, further studies are needed to test this hypothesis.

Not limited to GPVI collagen, the intracellular pathways of collagen degradation are also modulated by cytokines. This regulatory axis contributes to the inflammation-mediated injury/repair process in vitro (24). Il-1α, Il-1β, and Il-6, the early pro-inflammatory cytokines, decrease collagen synthesis by inducing MMP release/activity that degrades collagen (25, 26). Since uncleared collagen fragments are pro-inflammatory mediators, these cytokines regulate the inflammation-mediated response through removal of collagen fragments (24). Our analyses suggest that the GPVI collagen degradation is regulated by the activation of inflammatory cytokines-related pathways, namely Il-6 and wound healing signaling pathways. These pathways are affected by the upregulation of many major wound healing cytokines (Fig. 8 and 9A). These results suggest that the regulatory mechanisms at the early phase of PMDS involve the inflammation-mediated repair process via the transient upregulation of the wound healing SASP factors, leading to the GPVI collagen degradation.

Senescent cells accumulate around the wound, the tissue injury site, in vivo (mice and humans) (9, 27). This was shown by the significant increase of p21-, p16-, and SA-β-gal-positive cells in response to wounding (27–30). Those senescent cells contribute to optimal wound healing via SASP; the SASP mediates senescent cell clearance and tissue regeneration (9, 31). However, the origin of the senescent cells near the wound remained unclear. Here, we found that PMD induces the transient upregulation of wound healing SASP factors, accompanied by the transient increase of senescence markers, namely p53, p21, p16, and SA-β-gal, during PMDS. Hence, we speculate that PMD could be the physiological trigger of the senescent cell accumulation around the tissue injury sites.

Interestingly, the GPVI signaling pathway was also inhibited in CIS, associated with the transient upregulation of SASP factors namely *Il-6*, *Cxcl8*, *Mmp1*, and *Mmp3*. To date, no studies have reported that Ca^2+^ influx can lead to GPVI collagen degradation. Previous studies have shown that Ca^2+^ influx influences the cell morphology, proliferation, and collagen deposition of fibroblasts (32). Ca^2+^ influx increases the cell metabolic activity, migration, MMP production, collagen degradation, and cytokine release to accelerate the wound healing process (33). The Ca^2+^-dependent downregulation of the GPVI pathway can explain part of these phenotypes. In CIS, the transiently upregulated SASP factors, *Il-6* and *Cxcl*8, were not directly implicated in the wound healing signaling pathway but were implicated in Il-6 and Il-8 signaling pathways (Fig. S6 and S7A). The activation of Il-6 signaling also indirectly associated with the activation of Oncostatin M signaling to induce the transient upregulation of *Mmp1* and *Mmp*3. Oncostatin M is a multifunctional cytokine that belongs to the IL-6 subfamily (34).

It is known to promote ECM deposition and collagen degradation during the early inflammatory response in vitro (35, 36). These results suggest that the GPVI pathway was inhibited via different SASP-dependent pathways in PMDS and CIS.

Transient plasma membrane injury causes Ca^2+^ influx, which is essential for the membrane repair processes. The Ca^2+^-dependent membrane repair processes involve the toll-like receptors (TLR), NF-κB signaling, and pro-inflammatory cytokines (32). These mechanisms trigger the expression of stress response genes and promote wound healing (37). These results are consistent with our finding that Ca^2+^ influx contributes to the inhibition of the GPVI signaling pathway via the transient upregulation of SASP factors including the inflammatory cytokines and MMPs.

At the later phase of all senescence subtypes, *Il-6* and *Ccl2* were persistently upregulated. Our pathway comparison analyses indicate that the persistent upregulation of *Ccl2* led to the activation of HMGB1 signaling (Fig. 9). HMGB1 activates TLR signaling to induce the SASP production and accelerate paracrine senescence (38). Persistent DDR activity is required for the induction of several proinflammatory SASP factors, particularly IL-6 (39). Therefore, we speculate that the persistent upregulation of *Il-6* at the later phase might be induced by DDR. These factors were implicated in senescence-associated pathways, namely STING-mediated, cytokines-induced, and inflammation-mediated senescence (Fig. 9A and S7A). These results suggest that, at the later phase, all senescent cell subtypes tested here show senescent phenotypes at least partly mediated by the common SASP factors, *Il-6* and *Ccl2*.

## CONCLUSION

Plasma membrane damage induces transient upregulation of wound healing SASP factors, particularly *Il-6*, *Mmp1*, and *Mmp3,* accompanied by the increase of senescence marker proteins, p53, p21, and p16, in normal human fibroblasts. These results raise a possibility that the PMDS cells may contribute to wound healing in vivo. The upregulation of SASP in the PMDS cells could be triggered by Ca^2+^ influx, leading to the inhibition of GPVI collagen signaling (Fig. 11, left). The inhibition of GPVI collagen signaling further upregulates *Il-6*, *Mmp1*, and *Mmp3*, forming a positive feedback regulatory network. In the later phase of cellular senescence, *Ccl2* and *Il-6* are persistently upregulated in PMDS, CIS, and DDRS (Fig. 11, right). Thus, based on our findings, we propose that in early senescence, SASP mediates diverse cellular functions depending on senescent cell subtypes. In contrast, SASP becomes relatively uniform in late senescence, contributing to further deepening senescence.

**Fig. 11.**
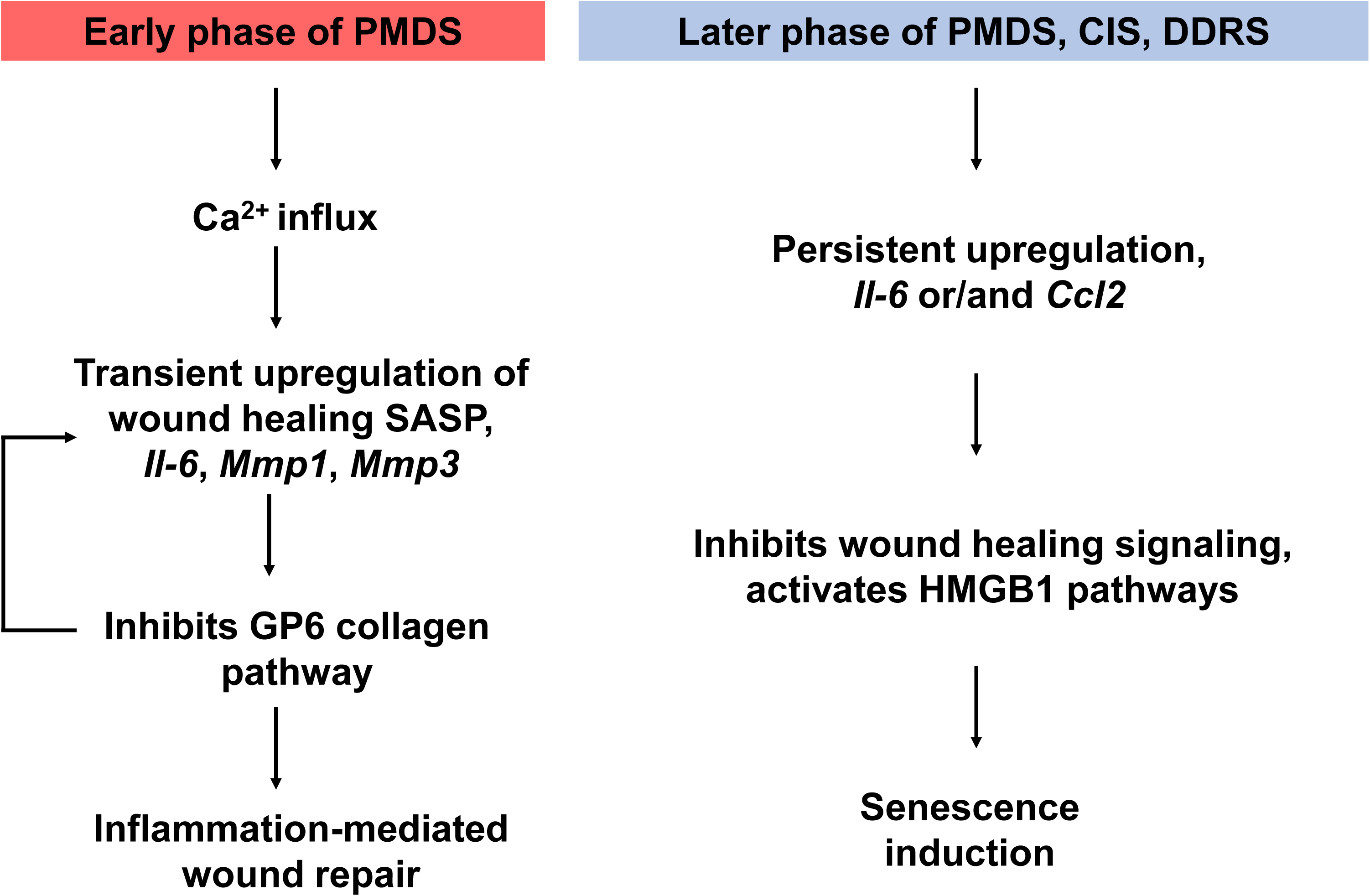
Schematic illustration shows the potential mechanisms regulating the SASP upregulation at the early phase of PMDS and the later phase of all senescence inductions.

## AVAILABILITY

Gene Ontology is an open-source collaborative tool available at (http://geneontology.org).

## ACKNOWLEDGEMENT

We are grateful to Barbee H. and Sugiyama S. for critical reading of the manuscript. We are grateful for the help and support provided by the Sequencing Section of the Research Support Division at the Okinawa Institute of Science and Technology Graduate University.

## TABLE AND FIGURES LEGENDS

Table S1. Primer sequences used for RT-qPCR analyses.

Table S2. SASP factors identified from the significant genes expressed in RS, PMDS, CIS, and DDRS compared to the SASP Atlas.

**Figure S1.**
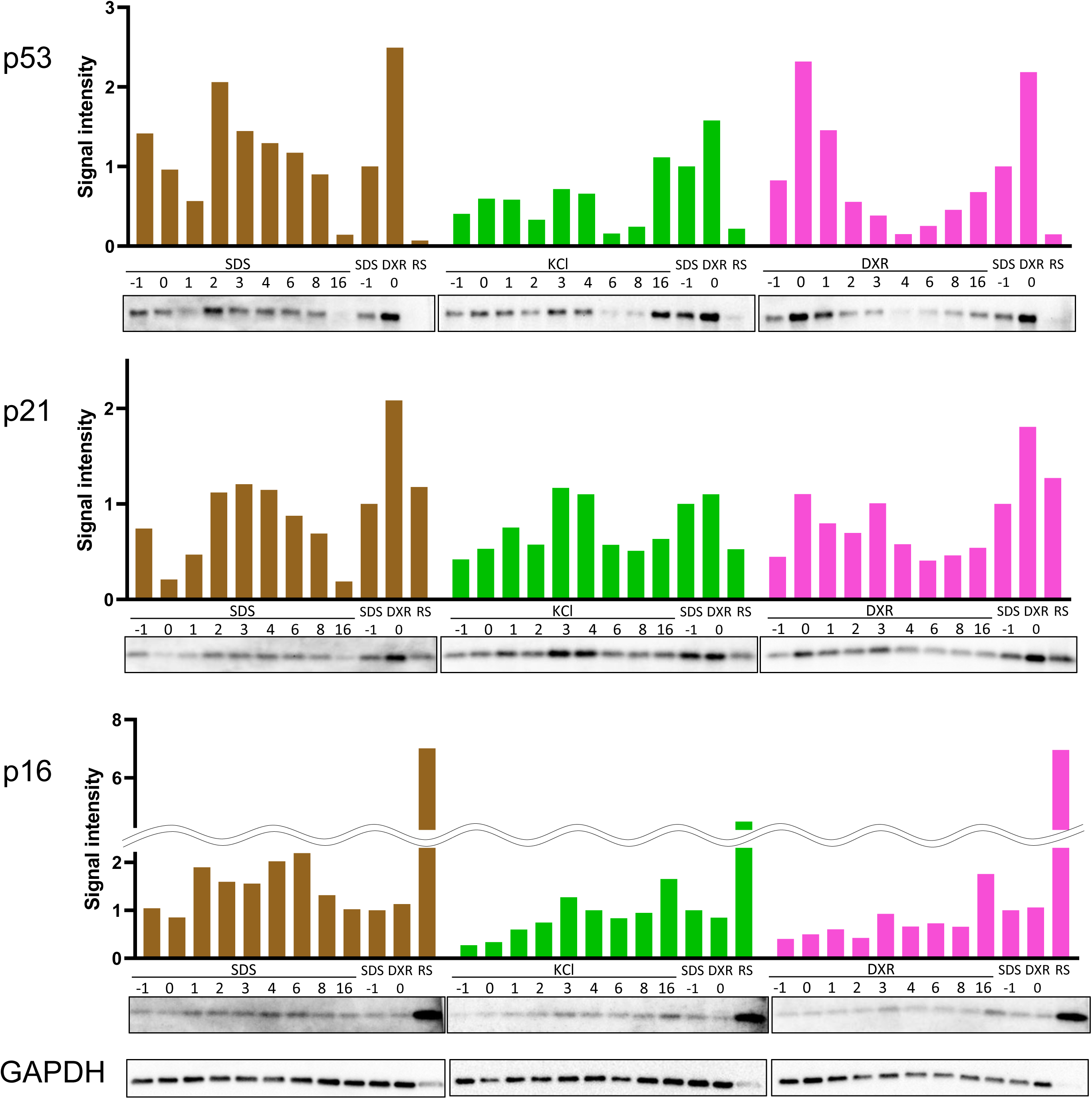
Quantification of Western blot. Graph shows the intensity of each band which corresponds to protein expression levels of either p53, p21, or p16. The protein expression levels were normalized by the intensity of SDS D-1 samples.

**Figure S2:**
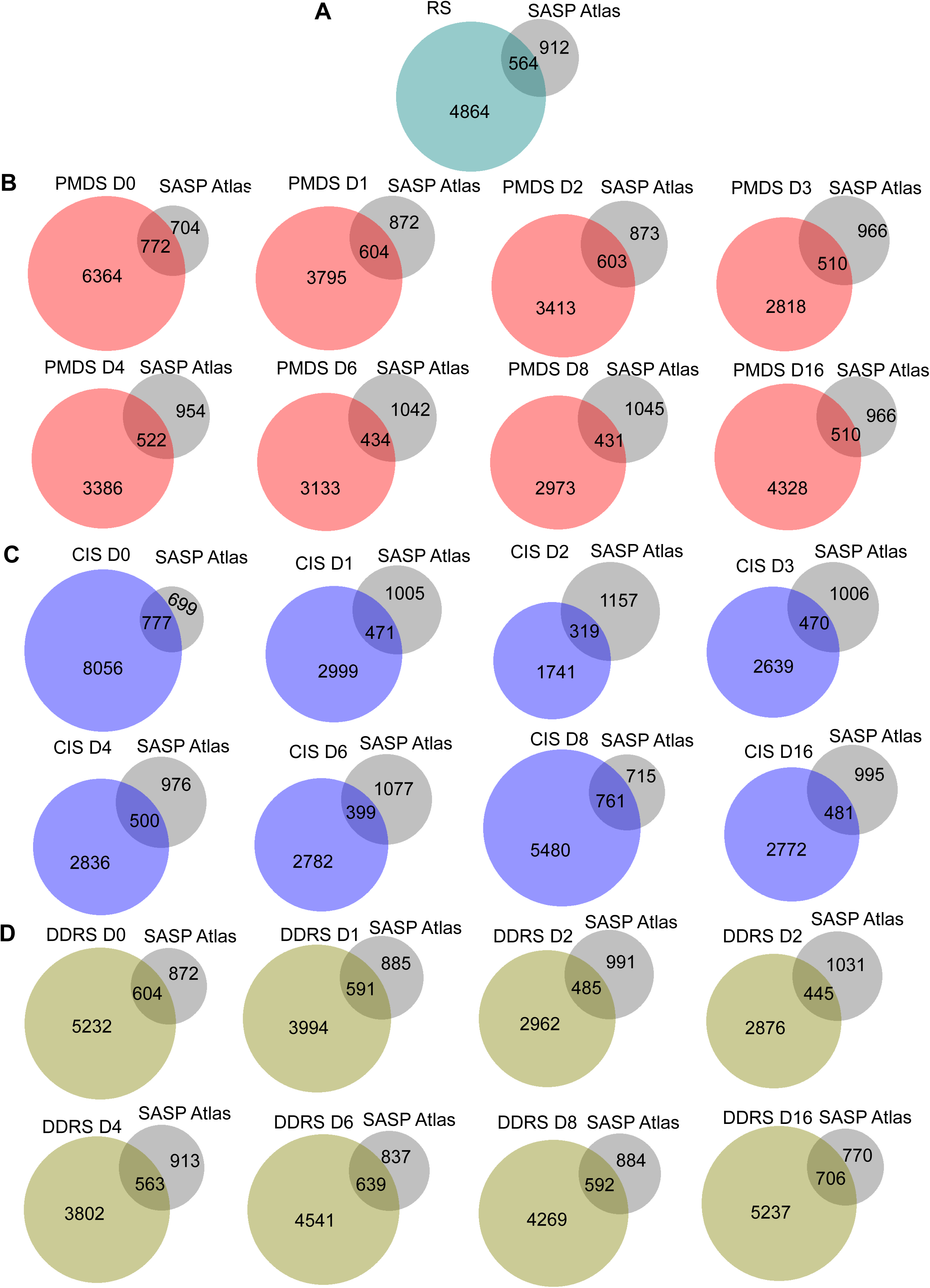
Venn diagram shows the overlapping SASP factors (intersections) in each time phase of RS (A), PMDS (B), CIS (C) and DDRS (D) with the SASP Atlas.

**Figure S3.**
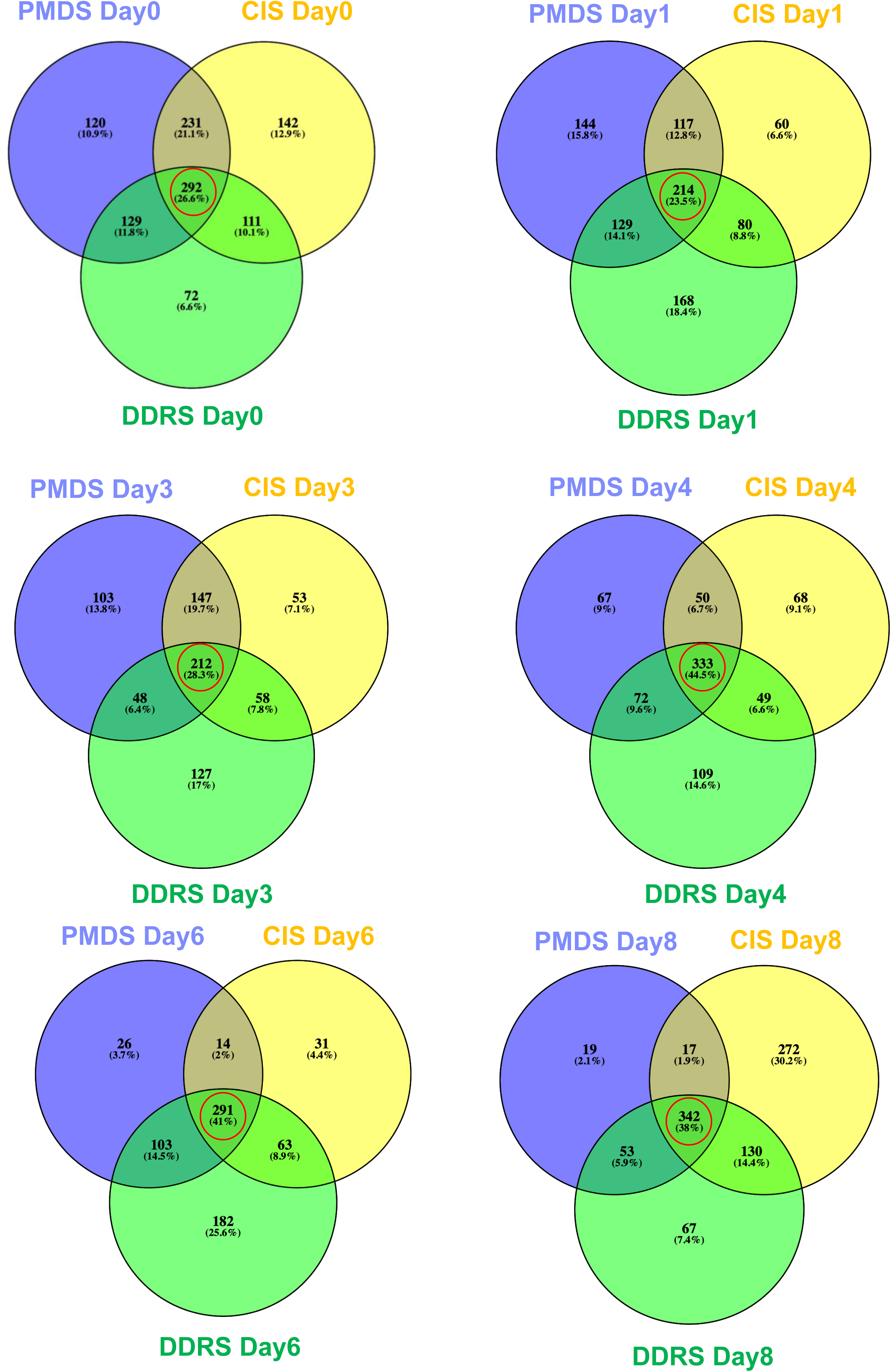
SASP factors are homogenous at the later phase of senescence induction. Venn diagram shows the overlapping genes (red circle intersections) in PMDS, CIS, and DDRS on Day 0, 1, 3, 4, 6 and 8. The intersection (red circle) shows the number of overlapping genes in all senescence inducers increased from the early phase (Day 0) to the later phase (Day 8).

**Figure S4.**
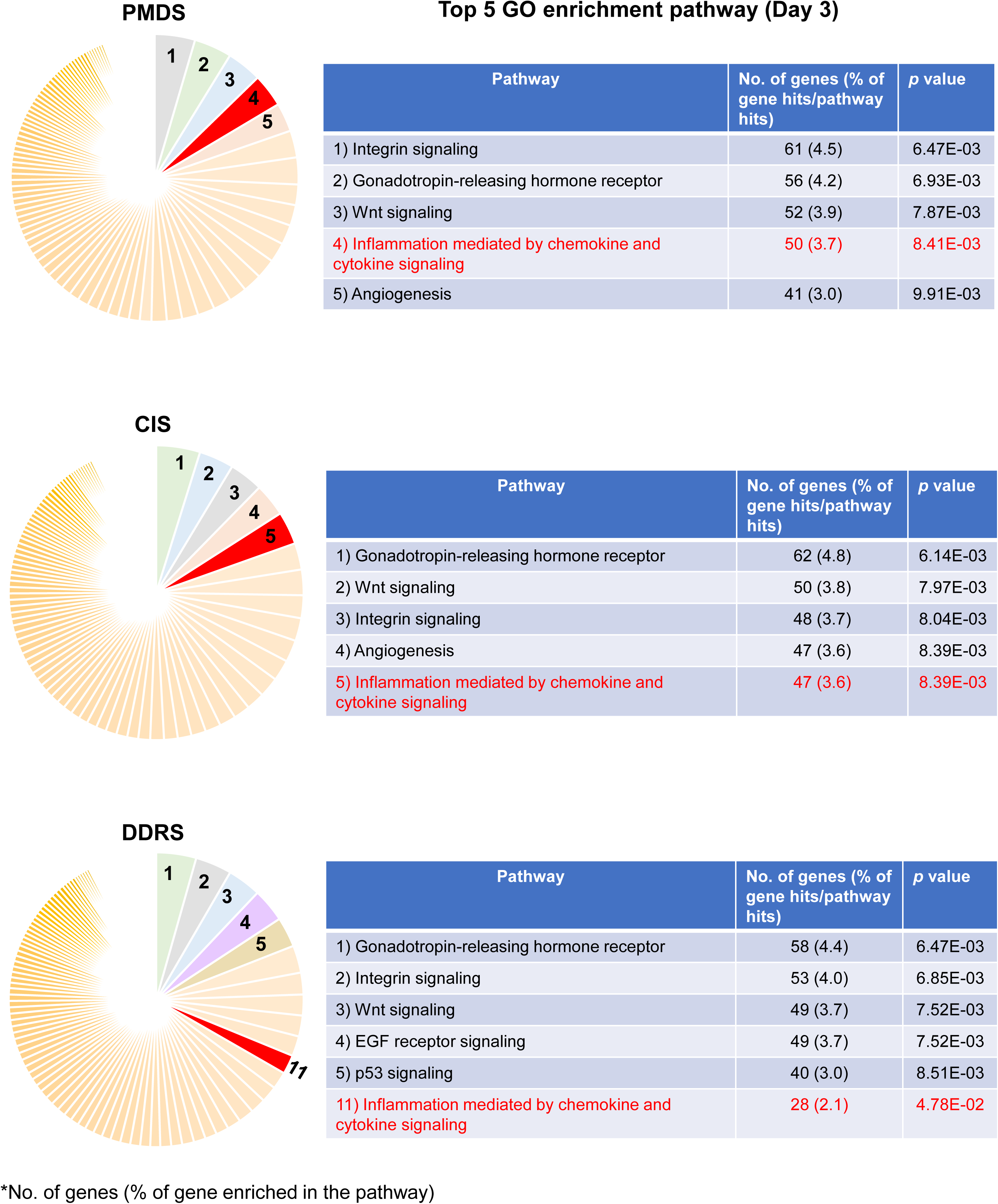
SASP-associated pathway is enriched only at the early phase of PMDS and CIS (Day 3). Pie chart and tables show the top five hit pathways for the enrichment of Gene Ontology (GO) SASP-associated pathway only in the early phase of PMDS and CIS (Day 3). The GO was analysed using PANTHER classification system (http://geneontology.org).

**Figure S5.**
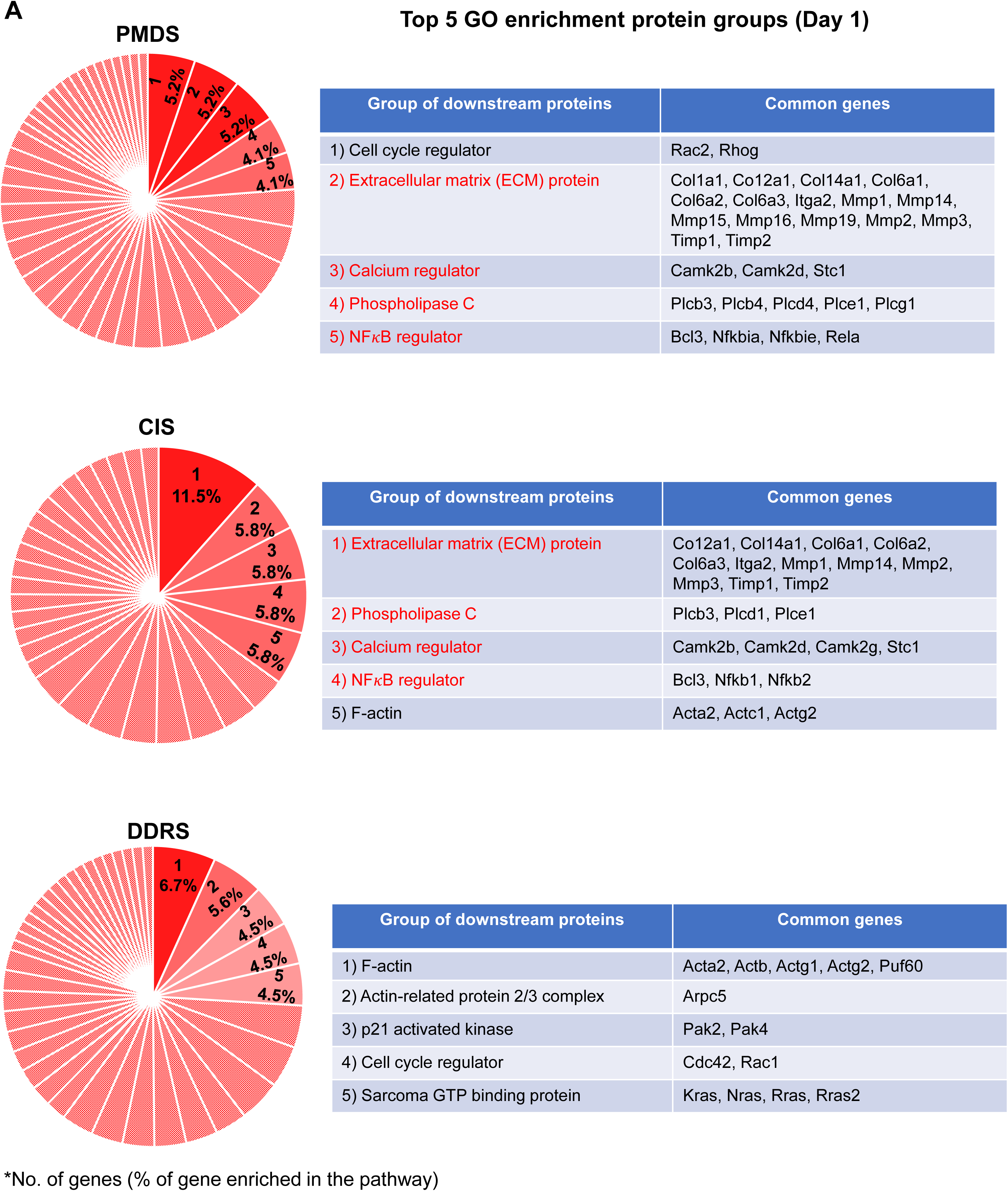
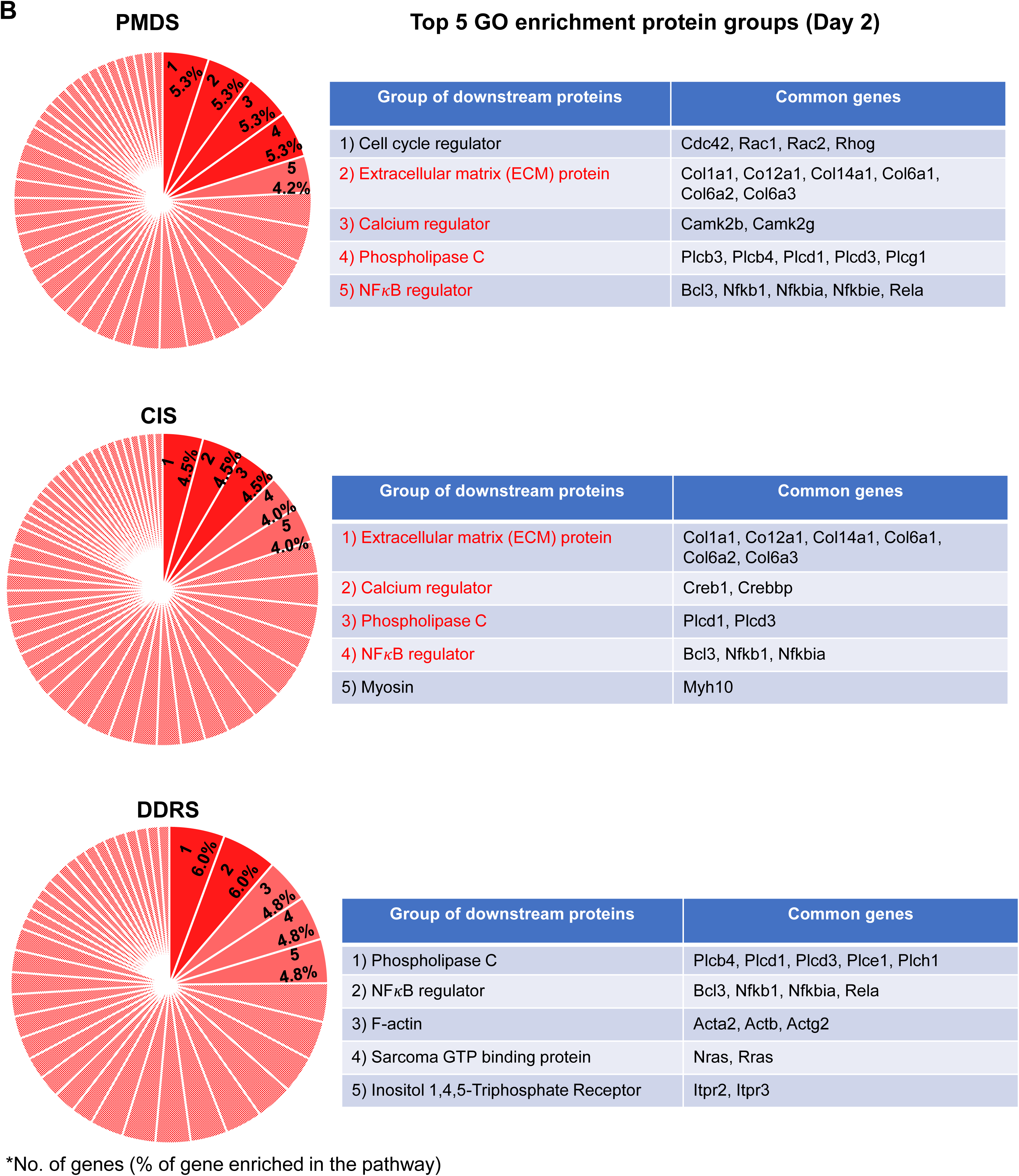
Similar downstream targets were identified in the SASP-associated pathways at the early phase of PMDS and CIS. Pie chart and tables show the top five hit categories for the enrichment of Gene Ontology (GO) for the sub-categories of downstream targets under the SASP-associated pathway at the early phase, Day 1 (A) and Day 2 (B) of PMDS, CIS, and DDRS. The GO was analysed using PANTHER classification system (http://geneontology.org).

**Figure S6.**
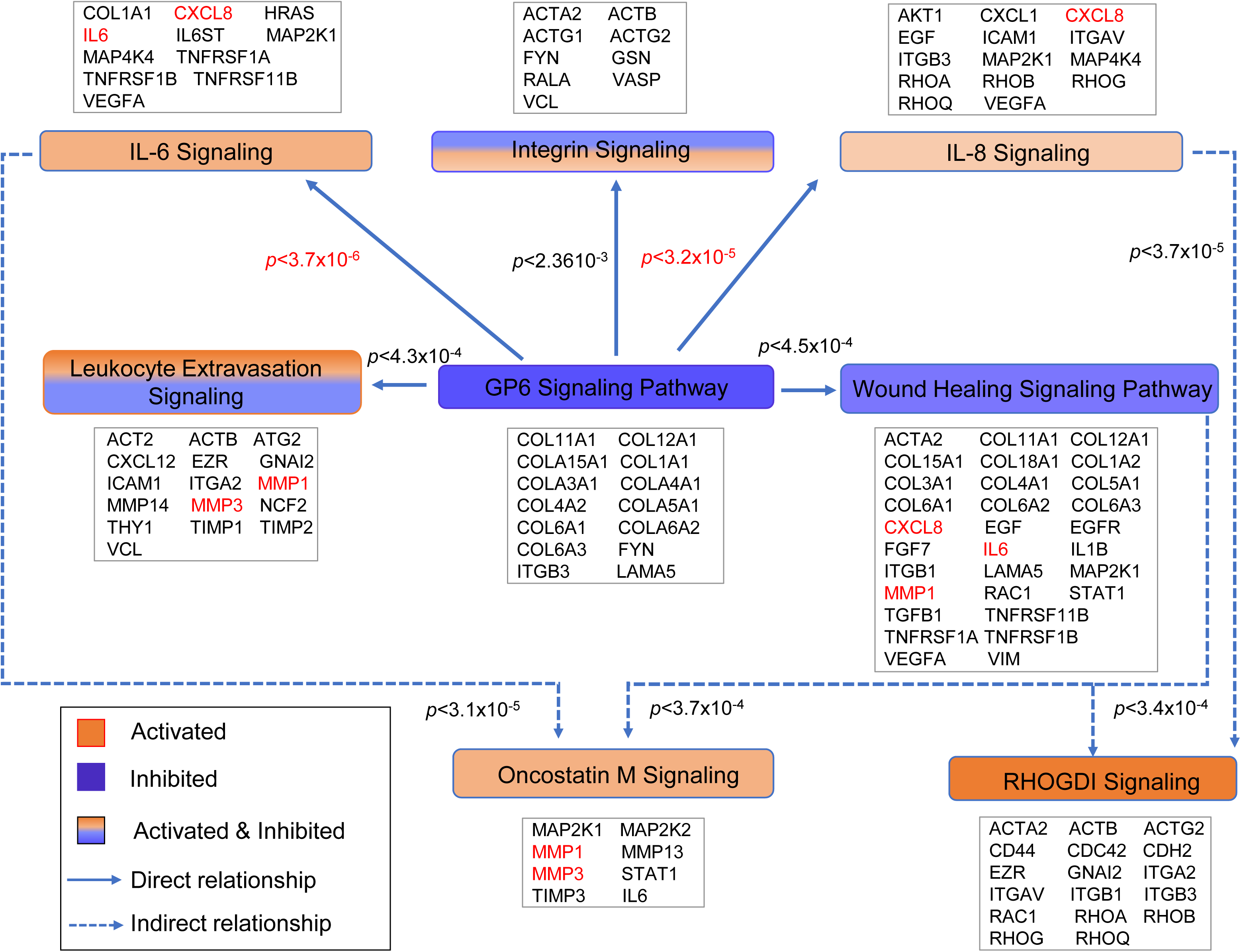
GPVI signaling pathways are implicated in common SASP factors regulated at the early phase of CIS, similar to PMDS. Diagram shows the significant GPVI signaling-associated pathways affected by CIS analysed by IPA software. Pie chart shows that common SASP factors, namely *Il-6*, *Mmp1*, and *Mmp3* (red fonts), are the downstream genes in most of the GPVI signaling-associated pathways. Orange and blue bars indicate positive and negative activation z-score respectively.

**Figure S7.**
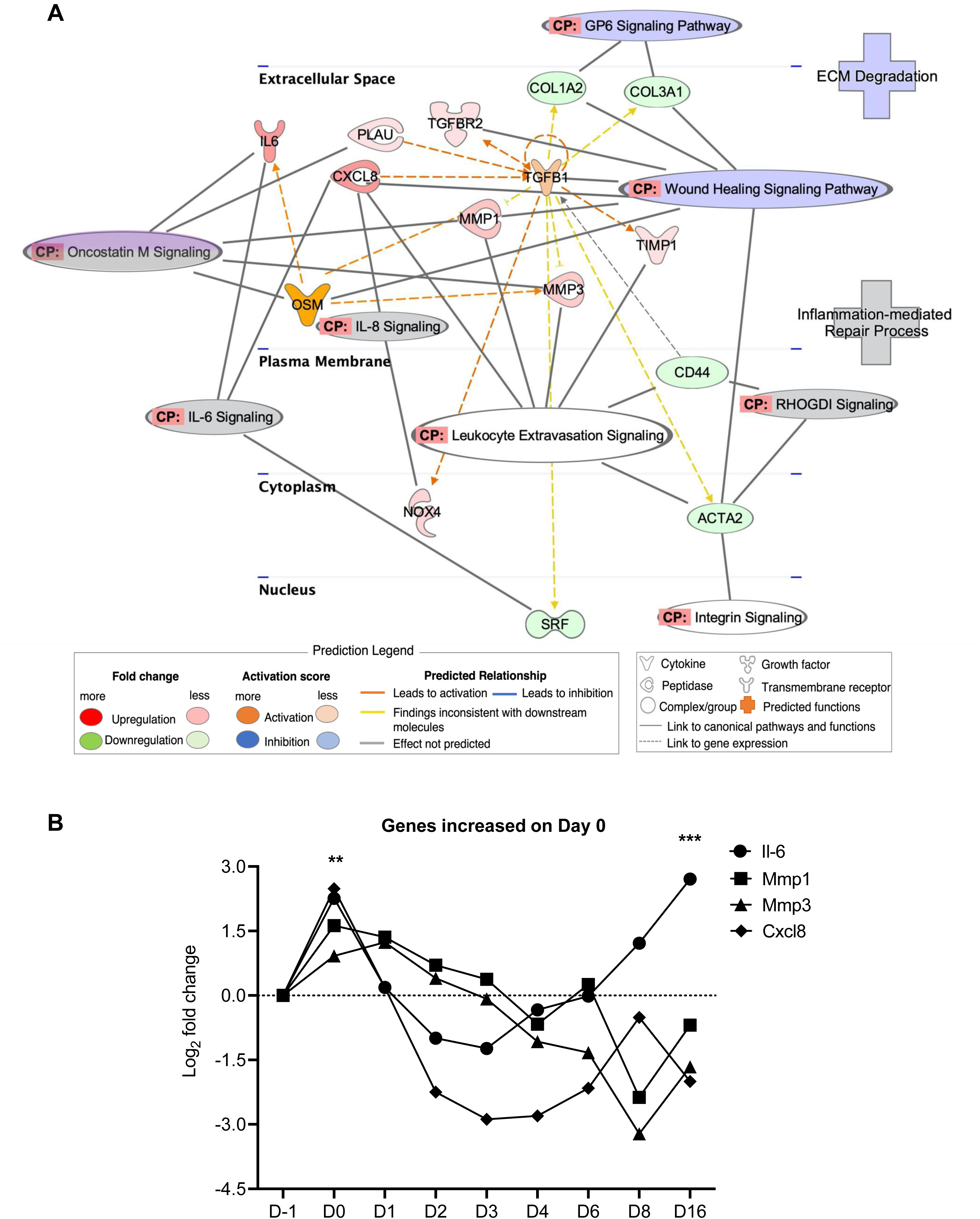
GPVI signaling is inhibited by transient increase of Il-6, Il-8, and Oncostatin M signaling-associated SASP factors early after CIS. IPA graphical representation shows the upstream regulators of GPVI signaling inhibition are mainly involved with Il-6, Il-8, and Oncostatin M signaling-associated SASP factors (A). The molecular interactions between the upstream regulators and the other SASP factors are displayed graphically as nodes (genes) and dotted lines (the biological activation or inhibition between the genes). Nodes in red indicate upregulated genes while those in green represent downregulated genes. Graph shows the expression of SASP factors regulated in CIS from Day 0-16. *Il-6, Cxcl8*, *Mmp1*, and *Mmp*3 were transiently upregulated at Day 0 (B).

**Figure S8.**
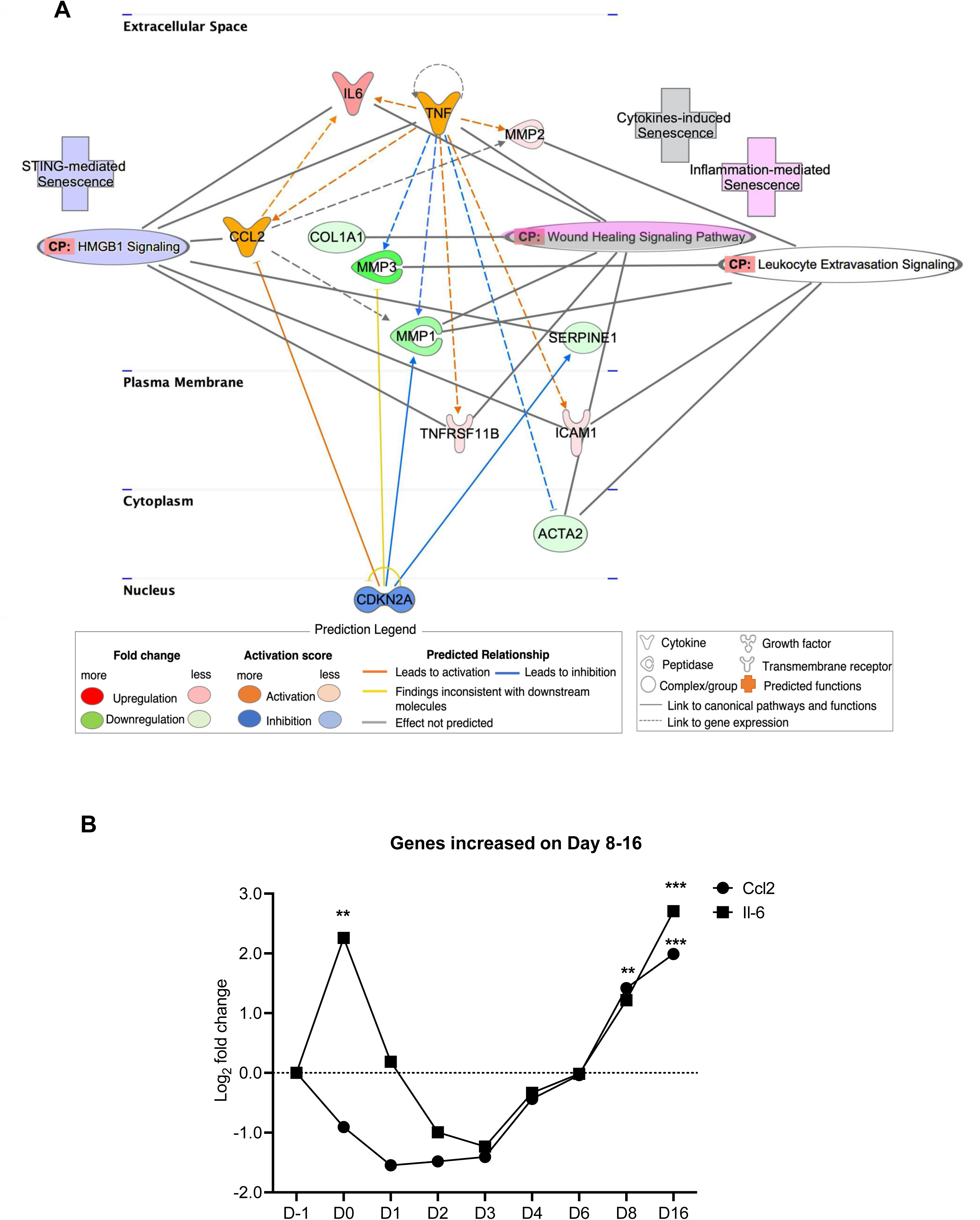
*Ccl2* and *Il-6* induce senescence late after CIS, comparable to PMDS and DDRS. IPA graphical representation shows *Ccl2* and *Il-*6 as the main upstream regulators, together with *Tnf* and *Cdkn2a*, in the later phase of PMDS and DDRS, which induce senescence, particularly by affecting HMGB1 and wound healing signaling pathways (A). The molecular interactions between the upstream regulators and the other SASP factors are displayed graphically as nodes (genes) and dotted lines (the biological activation or inhibition between the genes). Nodes in red indicate upregulated genes while those in green represent downregulated genes. Graph shows the expression of SASP factors, *Ccl2* and *Il-6*, regulated in CIS from Day 0-16. *Ccl2* and *Il-6* were persistently upregulated from Day 8-16 in CIS (B).

**Figure S9.**
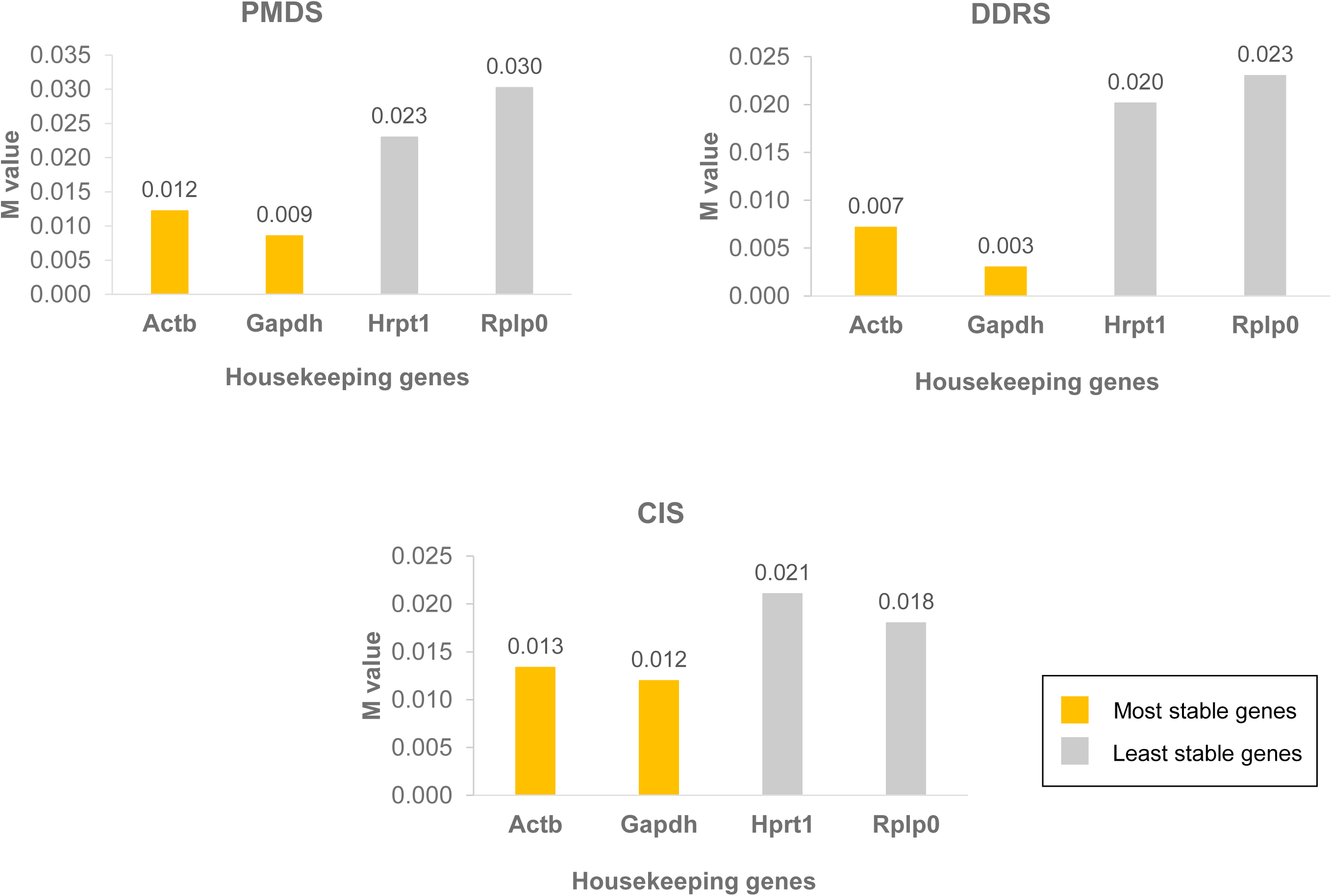
*Actb* and *Gapdh* are the ideal housekeeping genes for qPCR normalization. Bar graphs show the stability validation of the housekeeping genes, *Actb*, *Gapdh*, *Hprt1*, and *Rplp0*, analysed by the NormFinder software for the qPCR analyses of control, RS, PMDS, CIS, and DDRS. The most stable housekeeping gene has the lowest M value. NormFinder also calculates the stability value for the best combination of two genes.

**Figure S10.**
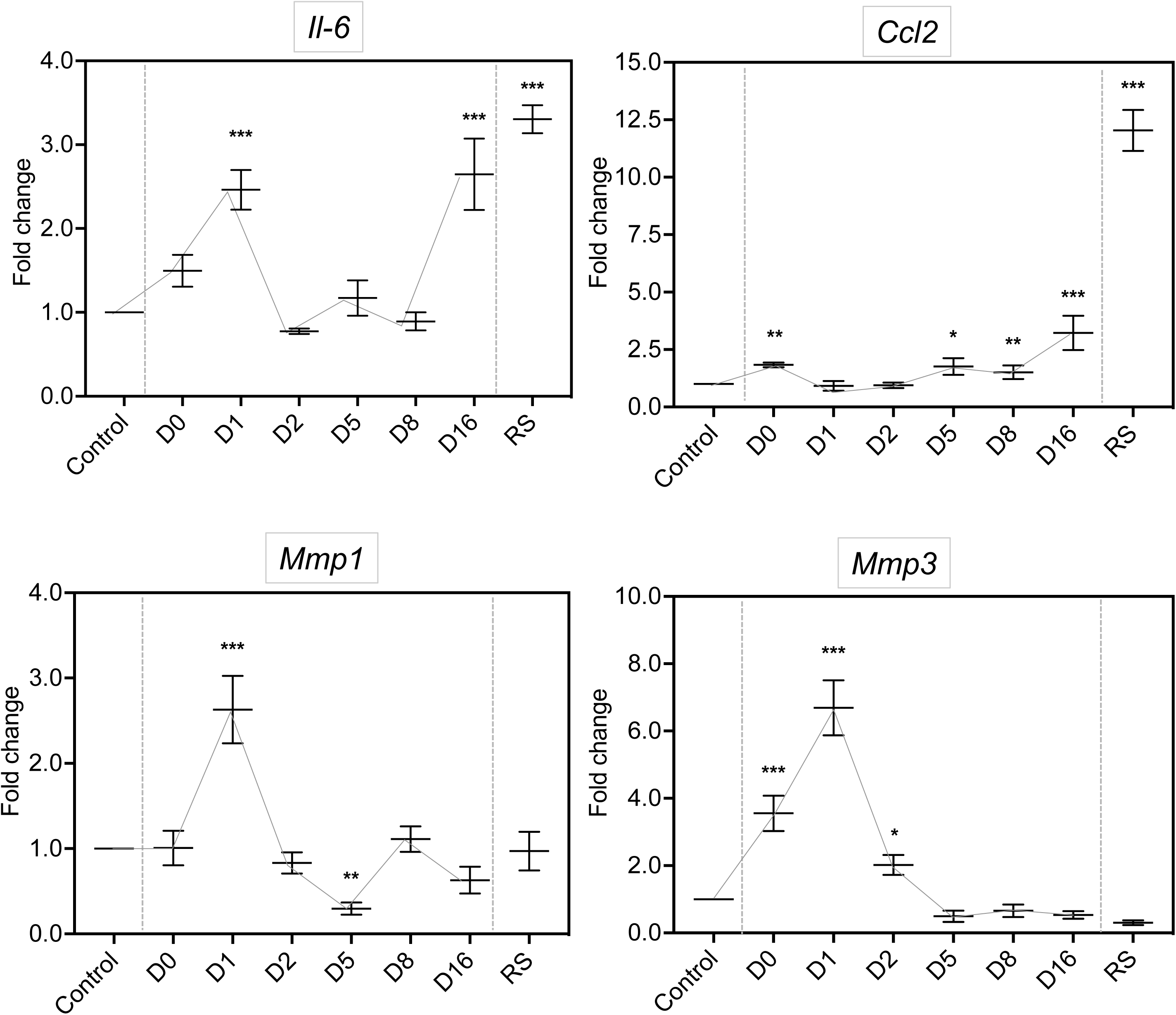
CIS SASP factors upregulation is confirmed by qPCR. Graph shows the mRNA expression of the SASP factors, *Il-6*, *Mmp1,* and *Mmp3*, which were transiently upregulated at Day 0-1 in CIS. Graph also shows the mRNA expression of SASP factors, *Ccl2* and *Il-6*, which were persistently upregulated from Day 5-16 in CIS. The expression of all SASP factors was normalized to that of *Actb* and *Gapdh* and calculated as fold changes. Error bars indicate standard deviation. *p<0.05, **p<0.01, ***p<0.001, compared to control, (n=3).

**Figure S11.**
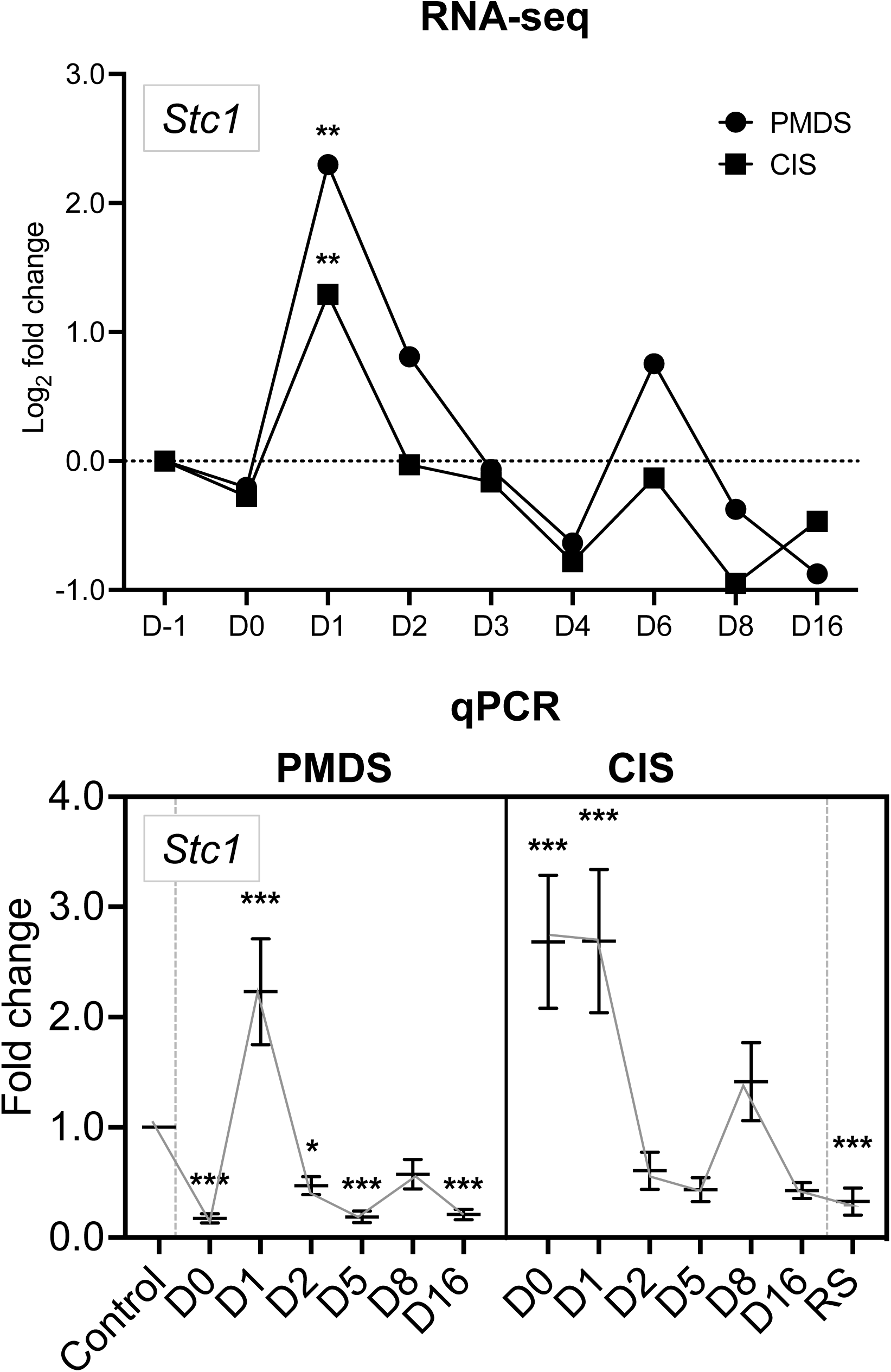
Transient upregulation of *Stc1* was observed early after PMDS and CIS. Graph shows the mRNA expression of the SASP factor, *Stc1,* which was transiently upregulated at Day 0-1 in CIS. A similar upregulation pattern was observed in both RNA-seq and qPCR analyses. The expression of all SASP factors was normalized to that of *Actb* and *Gapdh* and calculated as fold changes. Error bars indicate standard deviation. *p<0.05, **p<0.01, ***p<0.001, compared to control, (n=3).

