## Supplementary material for "Time-resolved transcriptomic profiling of senescence-associated secretory phenotype (SASP) in multiple senescent cell subtypes": Table S1

Table 1: Primer sequences used for RT-qPCR analyses

Gene	Forward primer	Reverse primer	Gene ID

__________________________________________________________________________________

*Actb*	5’-CATCACCATTGGCAATGAGC-3’	5’- TAGTTTCGTGGATGCCACAG -3’	60

*Gapdh*	5’-CGTCATGGGTGTGAACCATG-3’	5’-GTTGTCATGGATGACCTTGG-3’	2597

*Hprt1*	5’-TGCTGAGGATTTGGAAAGGG-3’	5’-GAGGGCTACAATGTGATGGC-3’	3251

*Rplp0*	5’-CTGGAAGTCCAACTACTTCC -3’	5’-GGATCTGCTGCATCTGCTTG-3’	6175

*Ccl2*	5’-AGGAAGATCTCAGTGCAGAG-3’	5’-GGTCAGCACAGATCTCCTTG-3’	6347

*Il-6*	5’-CCACTCACCTCTTCAGAACG-3’	5’-CCTCTTTGCTGCTTTCACA-3’	3569

*Mmp1*	5’-CTCAGTTTGTCCTCACTGAG-3’	5’-TTCTCAATGGCATGGTCCAC-3’	4312

*Mmp3*	5’-CAGGGATTAATGGAGATGCC-3’	5’-GAGTGGCCAATTTCATGAGC-3’	4314

*Stc1*	5’-ATGTGTGCAGCATCGCCAAG-3’	5’-ATCACATTCCAGCAGGCTTC-3’	6781
